# Dynamic ensemble of HIV-1 RRE stem IIB reveals non-native conformations that disrupt the Rev binding site

**DOI:** 10.1101/498907

**Authors:** Chia-Chieh Chu, Raphael Plangger, Christoph Kreutz, Hashim M. Al-Hashimi

## Abstract

The HIV-1 Rev response element (RRE) RNA element mediates the nuclear export of intron containing viral RNAs by forming an oligomeric complex with the viral protein Rev. Stem IIB and nearby stem II three-way junction nucleate oligomerization through cooperative binding of two Rev molecules. Conformational flexibility at this RRE region has been shown to be important for Rev binding. However, the nature of the flexibility has remained elusive. Here, using NMR relaxation dispersion, including a new strategy for directly observing transient conformational states in large RNAs, we find that stem IIB alone or when part of the larger RREII three-way junction robustly exists in dynamic equilibrium with non-native ‘excited state’ (ES) conformations that have a combined population of ~20%. The ESs disrupt the Rev binding site by changing local secondary structure and their stabilization via point substitution mutations decreases the binding affinity to the Rev arginine-rich motif (ARM) by 15- to 80-fold. The ensemble clarifies the conformational flexibility observed in stem IIB, reveals long-range conformational coupling between stem IIB and the three-way junction that may play roles in cooperative Rev binding, and also identifies non-native RRE conformational states as new targets for the development of anti-HIV therapeutics.

## INTRODUCTION

RNAs are growing in their importance as regulators of gene expression (1–3), novel drug targets (4–6), and as tools for bioengineering applications and synthetic biology (7,8). RNA and other biomolecules do not fold into a single structure but rather form a statistical ensemble of many interconverting conformations (9–12). The biological activities of nearly all non-coding RNAs depend on these dynamics. For example, binding of ligands, proteins, other RNAs, or changes in physiological conditions such as temperature can favor specific conformations in the ensemble and cause a structure specific change in activity (13–15). Thus, a deep understanding of how RNAs function within cells requires an understanding of their dynamic behavior. This in turn may enable the targeting of RNA in drug discovery efforts (12,13) as well as the rational design of RNA-based devices (7).

Application of different biophysical techniques have led to certain themes that reoccur in the ensemble description of RNA (15). One theme is that most if not all RNAs (other than A-form duplexes) transiently form alternative conformations *in vitro* and *in vivo* that feature non-native secondary structure (9,16–20). These alternative conformations, often referred to as ‘excited states’ (ESs) (16,17,21), form on the micro- to-millisecond timescale and involve small differences in base-pairing in and around non-canonical motifs such as bulges and internal loops. These dynamic transitions can serve as facile switches (22,23) or help break down larger conformational transitions into multiple kinetically labile microscopic steps (18,24,25). Given the growing observation of such non-native conformations in a variety of RNAs by NMR (26–28) and chemical probing (29–31), it is important to examine their biological roles, their potential as new drug targets, and to assess how they might impact the interpretation of mutagenesis and structure probing *in vitro* and *in vivo*.

The Rev response element (RRE) from the human immunodeficiency virus type 1 (HIV-1) (32–35) is an example of a flexible RNA drug target (36–41) has the potential to adopt such alternative non-native conformations (Fig. 1A) (42–44). RRE is a ~350 nucleotide *cis*-acting RNA element located within the *env* gene (32–34). In HIV-1, RRE mediates export of unspliced or partially spliced viral RNAs to the cytoplasm by coordinating the assembly of multiple molecules of the viral protein Rev to form a homo-oligomeric ribonucleoprotein complex (35,45–48). The assembly of this complex has been the subject of many investigations (42,49–53). RRE folds into a structured RNA with five distinct stem loops (I to V) (46,54,55). Studies indicate that two Rev molecules bind cooperatively to high affinity sites in stem IIB and the nearby stem II three-way junction (42,48,49,56,57) (Fig. 1A). This initial binding event is thought to nucleate assembly, which then extends to stem I, including a high affinity site in stem IA, ultimately resulting in the coordinated and sequential binding of multiple Rev copies through a series of Rev-Rev and Rev-RRE interactions (35,42,48,52). The high concentration of Rev required for assembly is thought to set an expression threshold such that Rev would only function during later stages of viral replication (35,45).

**Figure 1.**
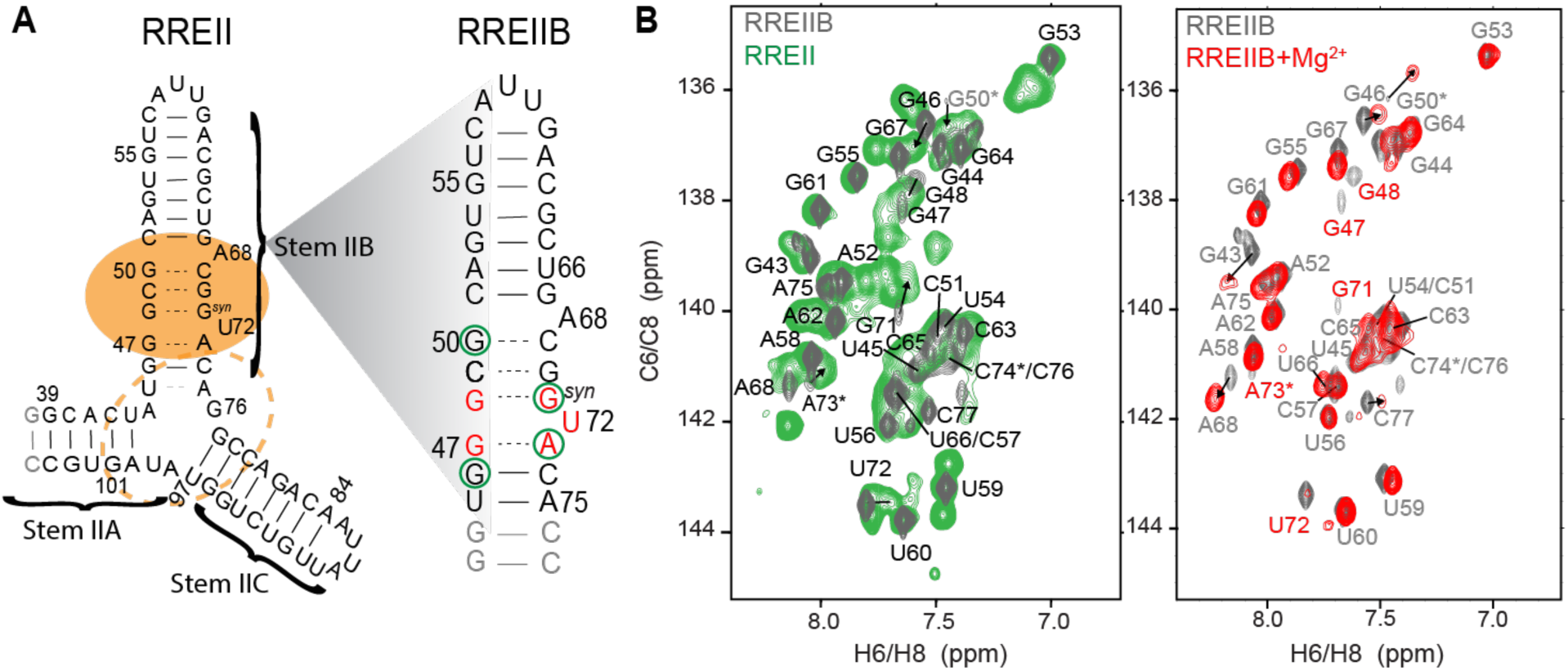
RREII constructs used in NMR studies. (A) Three-way junction (RREII) and stem IIB (RREIIB) constructs used in this study. The Rev primary and secondary binding sites are indicated using filled and dashed orange circles, respectively. Base pairs with and without detectable H-bonds are indicated using solid and dashed lines, respectively. Residues showing differences in chemical shifts between RREII and RREIIB or that have broadened resonances indicative of micro-to-millisecond exchange are highlighted using green circles and colored red on RREIIB, respectively. (B) Comparison of aromatic 2D HSQC of RREIIB (black) with RREII (green) and RREIIB plus Mg^2+^ (red). Buffer conditions: 15 mM sodium phosphate, 25 mM NaCl, 0.1 mM EDTA, pH 6.4 with or without 3 mM MgCl_2_. Resonances exhibiting line-broadening are labeled in red and those with ambiguous assignments denoted using an asterisk.

Because of its central importance in viral mRNA export and HIV replication (32,58–61), stem IIB has been the subject of numerous biochemical and structural studies including efforts to target RRE in the development of anti-HIV therapeutics (36–40). NMR and X-ray structures of isolated stem IIB in complex with Rev arginine-rich motif (ARM) show that RNA recognition leads to conformational changes that widens the major groove to accommodate binding of ARM as an α-helix that forms base-specific hydrogen bonds and electrostatic interactions with backbone phosphate groups (43,44,56,62–66). Biochemical studies show that an additional Rev molecule binds cooperatively at the nearby stem II three-way junction (45,48,67). A recent crystal structure of stem IIB containing an engineered junction site with a Rev dimer shows that the junction site helps orient the Rev subunits through interactions that depend on the junction architecture (56). Biochemical studies indicate that flexibility at the junction is important for adaptive recognition of the second Rev molecule through non-specific interactions and is also an important determinant of binding affinity and cooperativity (48,49,68). Indeed, the high SHAPE reactivity observed for nucleotides within stem IIB and the stem II three-way junction within the larger ~234 and ~354 nt RRE contexts indicates that this region is highly flexible. NMR studies of isolated stem IIB also provide evidence for flexibility in the purine rich region (43,44). However, the nature of this flexibility has remained elusive.

In this study, we used NMR relaxation dispersion (RD) techniques to examine the dynamics of RRE stem IIB in isolation and when part of the three-way junction. The data indicate that stem IIB forms a dynamic ensemble of three conformations including two non-native ESs that remodel key structural elements required for Rev binding. We discuss potential roles for the ESs in cooperative Rev binding and propose that they provide new targets for the development of anti-HIV therapeutics.

## MATERIALS AND METHODS

### Preparation of RNA samples

RREIIB and RREII were prepared by *in vitro* transcription using T7 RNA polymerase (New England Biolabs Inc.), chemically synthesized DNA templates (Integrated DNA Technologies) with designed T7 promoter sequence (TTAATACGACTCACTATA) and uniformly ^13^C/^15^N-labeled nucleotides triphosphates (Cambridge Isotope Laboratories Inc.). RREIIB mutants (RRE20, G48A/G50U-RREIIB, A68C-RREIIB, U72C-RREIIB, A68C/G50A/C69U-RREIIB, and m^3^U72-RREIIB), and RREII mutants (A68C-RREII, U72C-RREII, A68C/G50A/C69U-RREII) were synthesized using an in-house oligo synthesizer (MerMade 6, BioAutomation) with solid-phase RNA synthesis using N-acetyl protected 2’-tBDSilyl-phosphoramidites (ChemGenes Corporation) and 1 μmol standard columns (1000 Å, BioAutomation) with 4,4’-dimethoxytrityl (DMT)-off synthesis followed by base and 2’-O deprotection (100 μmol DMSO, 125 μL TEA•3FH), and ethanol precipitation. The DMT-off 2’-O deprotection is recommended for larger RNA synthesis to obtain cleaner NMR spectra.

A similar approach was used to prepare site labeled RREII (^15^N3-U72-RREII, ^13^C6-U72-RREII and ^13^C8-G71-RREII) using ^13^C/^15^N-atom-specifically-labeled RNA phosphoramidites (69). All RNA samples were purified using 20% (w/v) denaturing polyacrylamide (29:1) gel within 8M urea, 20 mM Tris Borate and 1 mM ethylene-diaminetetraacetate (EDTA) TBE buffer followed by Elutrap electro-elution system (Whatmann, GE healthcare) with 40 mM Tris Acetate and 1 mM EDTA (TAE) buffer then ethanol precipitation. The RNA pellets were dissolved in water and annealed by heating at 95 °C for 10 mins then rapidly cooling on ice. After measuring the concentration, the RNA samples were concentrated and buffer-exchanged three times into NMR buffer (15 mM sodium phosphate, 25 mM NaCl, 0.1 mM EDTA, with or without 3 mM MgCI_2_ at pH=6.4) using Amicon Ultra centrifugal filters (EMD Millipore). For NMR samples in 100% D_2_O, the RNA samples were flash frozen and lyophilized overnight before dissolving in 100% D_2_O (EMD Millipore).

### NMR experiments

#### Resonance assignment

NMR experiments were performed on Bruker Avance III 600-MHz or 700 MHz-NMR spectrometers equipped with 5 mm triple-resonance cryogenic probe at 10 or 25 °C. The RREIIB resonances in two-dimensional (2D) ^1^H-^15^N (N1/N3-H1/H3) and ^1^H-^13^C (C2/C6/C8-H2/H6/H8) heteronuclear single quantum coherence (HSQC) spectra were assigned using 2D ^1^H-^1^H nuclear overhauser effect spectroscopy (NOESY) with 180 ms mixing time for exchangeable proton at 10 °C with 10% D_2_O and 220 ms mixing time for non-exchangeable proton at 37 °C in 100% D_2_O in the absence of Mg^2+^. The same HSQC and NOESY (mixing time 180 ms) experiments were used to assign resonances in RRE20 and G48A/G50U RRE mutants at 25 °C in the absence of Mg^2^+. All NMR data were analyzed using NMRPipe (70) and SPARKY (T.D. Goddard and D.G. Kneller, SPARKY 3, University of California, San Francisco).

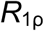 measurements. ^13^C RD experiments (71,72) were carried out on Bruker Avance III 700-MHz NMR spectrometer with 5-mm triple-resonance cryogenic probe at 25 °C using 1.0 mM ^13^C/^15^N-uniformly labeled RNA samples in NMR buffer (15 mM sodium phosphate, 25 mM NaCl, 0.1 mM EDTA, pH 6.4 with or without 3 mM Mg^2+^). On- and off-resonance RD data on aromatic (C8, C6) spins were measured with varying spinlock powers (*ω*2*π*^−1^ Hz) and off-sets (*Ω*2*π*^−1^ Hz) (Table S3) using seven delay times between 0 and 60 ms (73). Peak intensities for each delay time point were obtained using NMRPipe and fit to a monoexponential decay function using in-house python script to calculate 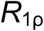 values (74). Errors in 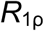 were estimated using Monte Carlo simulations with 500 iterations as previously described (75).

#### 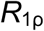 data analysis

The RD data was analyzed to obtain exchange parameters through numerical solutions of the Bloch-McConnell equations (21) using an in-house python script (23). The data was fit using two-state (individual or global) or three-state (individual or global) models with or without minor exchange using average effective field alignment as previously described (17,74). Global fitting of the data was carried out by sharing the populations and k_ex_ of ES1 and ES2 (Table S1). Model selection was preformed as previously described (23) using the Akaike’s (*w*_AIC_) and Bayesian information criterion (*w*_BIC_) weights to select the model with the highest relative probability (76,77).

### Fluorescence anisotropy binding experiments

Binding experiment were preformed using a Rev-ARM peptide labeled with 3’-end fluorescein (Rev-Fl, TRQARRNRRRRW RERQRAAAACK-FITC, LifeTein LLC) (78). Fluorescence anisotropy measurements were performed using a CLARIOstar plate reader (BMG LABTECH) using 480 nm excitation and 540 nm emission filter. A constant concentration of Rev-Fl (1 nM for RREII and RREII mutants, 10 nM for other constructs) was added into 384-well plate with serially diluted RNA in the reaction buffer (30 mM HEPES, pH= 7.0, 100 mM KCl, 10mM sodium phosphate, 10 mM ammonium acetate, 10 mM guanidinium chloride, 2 mM MgCl_2_, 20 mM NaCl, 0.5 mM EDTA, and 0.001% (v/v) Triton-X100) (78). K_d_ values were obtained by fitting the measured fluorescence anisotropy values to one-site binding equations using least-squares methods implemented in Mathematica 10.0 (Wolfram Research).

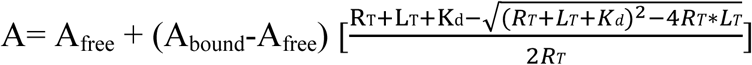

A is the measured value of fluorescence anisotropy; R_T_ is the total RNA concentration; L_T_ is the total Rev-Fl concentration; A_free_ is the anisotropy without Rev-Fl binding; A_bound_ is the anisotropy with saturated Rev-Fl binding; and K_d_ is the dissociation constant. The uncertainty in the fluorescence anisotropy was obtained based on the standard deviation over triplicate measurements.

## RESULTS

### Mg^2+^ and the three-way junction impacts the structural dynamics of stem IIB

Prior NMR studies were conducted in the absence of Mg^2+^ and focused exclusively on isolated stem IIB in which the stem II three-way junction was omitted and replaced with stable base pairs (43,44,62–64). However, considering the close proximity of stem IIB to the three-way junction, it is possible that such isolated constructs of stem IIB in the absence of Mg^2+^ do not fully capture the structural dynamics in the biologically relevant three-way junction context in the presence of Mg^2+^. We therefore used NMR to compare an isolated 35 nt construct of stem IIB (RREIIB) with a 68 nt construct (RREII) containing the stem II three-way junction (Fig. 1A). Both constructs avoid mutations used in prior NMR and X-ray studies of RREIIB (43,44,56,65,66).

2D CH HSQC spectra of RREIIB in the absence of Mg^2+^ revealed that many resonances in the internal loop (G47 to C51 and A68 to A73) are significantly broadened (Fig. 1B and S1). This was the first hint that stem IIB undergoes micro-to-millisecond timescale exchange with alternative ES conformations. Addition of 3 mM Mg^2+^ resulted in small chemical shift perturbations indicating minor changes in the dominant ground state (GS) structure (Fig. 1C and S1). In addition, Mg^2+^ further broadened resonances belonging to nucleotides in the purine-rich internal loop (G47, G48, G71, U72, A73), indicating increased micro-to-millisecond conformational exchange, whereas it reduced the broadening at the A68 bulge, indicating a decrease in micro-to-millisecond exchange (Fig. 1C and S2). Analysis of NOE distance-based connectivity and chemical shifts measured in RREIIB suggest a dominant GS conformation similar to that reported in prior NMR studies (43,44) with G48(*anti*)-G71(*syn*) and G47(*anti*)-A73(*anti*) mismatches, partially stacked A68 bulge, and flipped out U72 bulge (Fig. 1B and S1).

We find very good agreement when overlaying spectra of RREIIB with those of the RREII three-way junction both in the absence or presence of Mg^2+^ (Fig. 1B and S1). However, small differences are observed for G46 near the three-way junction and more distant residues such as G71 and G50 (Fig. 1B and S1B). In addition, the line broadening at the purine rich region was more severe in RREII as compared to RREIIB in both the absence and presence of Mg^2+^ (Fig. S2B). These results indicate that the three-way junction and Mg^2+^ have small effects on the GS conformation but do alter micro-to-millisecond conformational exchange at the internal loop region.

### Characterizing conformational exchange using NMR relaxation dispersion

To gain further insights into the conformational exchange, we carried out using 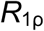 RD NMR experiments on RREIIB both in the presence and absence of Mg^2+^. The 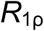 experiment (71,79) measures the chemical exchange contribution (*R*_ex_) to intrinsic transverse relaxation rate (*R*_2,int_) of NMR resonances during a relaxation period when the spin is irradiated with an applied radiofrequency pulse with varying powers (*ω*2π^−1^) and frequencies (*Ω*2*π*^−1^).

In the absence of Mg^2+^, RD consistent with micro-to-millisecond exchange was observed for most residues in the Rev binding site (G46-C8, G47-C8, G67-C8, A68-C8, G71-C8, U72-C6) (Fig. 2A). In contrast, no RD was observed for residues outside this region (A52-C8, G53-C8, U66-C6 and A75-C8) (Fig. 2A). Similar nucleotide specific RD was observed in the presence of 3 mM Mg^2+^ though the lower quality of the NMR spectra did not permit RD measurements for all residues (Fig. S2). However, the RD profiles differed in Mg^2+^ and were enhanced for U72-C6 while they were diminished for A68-C8 (Fig. 2B), consistent with the Mg^2+^ induced changes in line-broadening observed in 2D HSQC spectra (Fig. S2).

**Figure 2.**
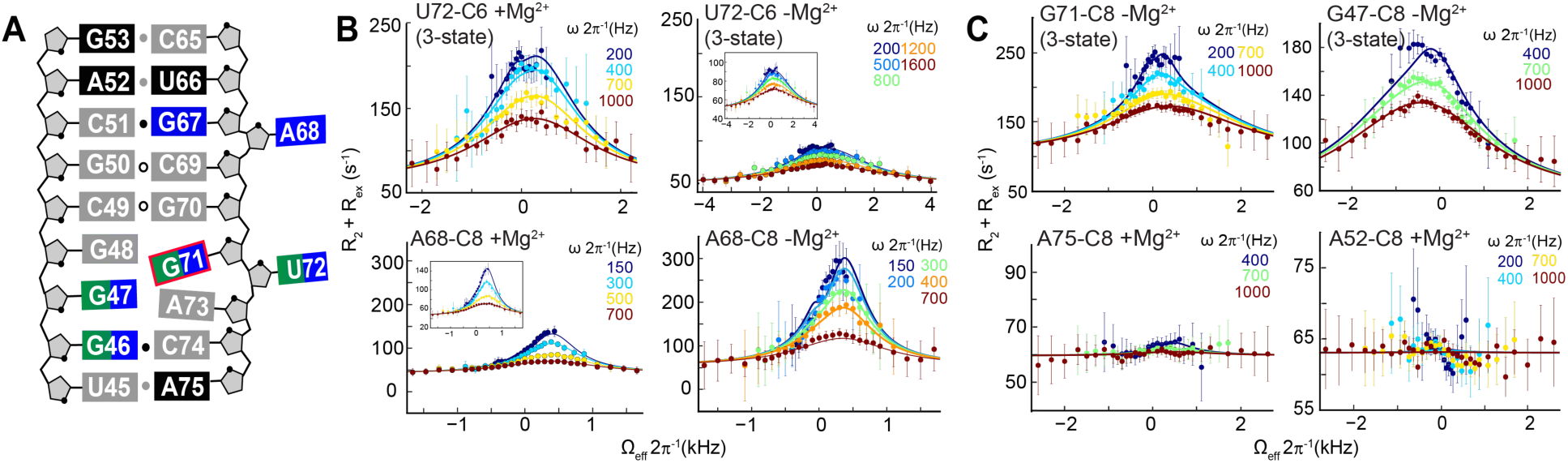
Off-resonance 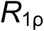 NMR measurements in RREIIB. (A) Secondary structure of RREIIB. Nucleotides with no RD, ES1, and/or ES2-specific RD are colored black, green and blue respectively. Nucleotides for which RD could not be measured are in gray. (B) Comparison of off-resonance 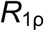 profiles in the presence and absence of 3 mM Mg^2^+. Different power levels (ω2π^_1^) are color-coded. Solid lines represent the global fits to the RD data. Error bars represent experimental uncertainty based on Monte Carlo analysis of monoexponential decay curves and the signal noise. (C) Representative examples of off-resonance 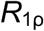 profiles showing resonances with three-state RD and absence of RD. Buffer conditions: 15 mM sodium phosphate, 25 mM NaCl, 0.1 mM EDTA, pH 6.4 with or without 3 mM MgCl_2_.

Fitting of the RD data (see methods) allowed determination of the ES population (p_B_), the exchange rate (k_ex_ = k_1_ + k_−1_), and the carbon chemical shift differences between the GS and ES (Δω = ω_ES_ − ω_GS_), which carry information regarding the ES conformation (80). Most RD data could be satisfactorily fit to a two-state (GS ⇌ ES) model (Fig. 2B, 2C and S3A). However, for U72-C6, G71-C8, G46-C8 and G47-C8 (Fig. 2B, 2C and S3A), the asymmetric RD profiles called for a three-state (ES1 ⇌ GS ⇌ ES2) fit with star-like topology (Fig. 3A) (17,80). The RD data therefore indicate that in RREIIB, stem IIB transiently forms at least two distinct ES conformations both in the presence and absence of Mg^2+^ (Table S1).

**Figure 3.**
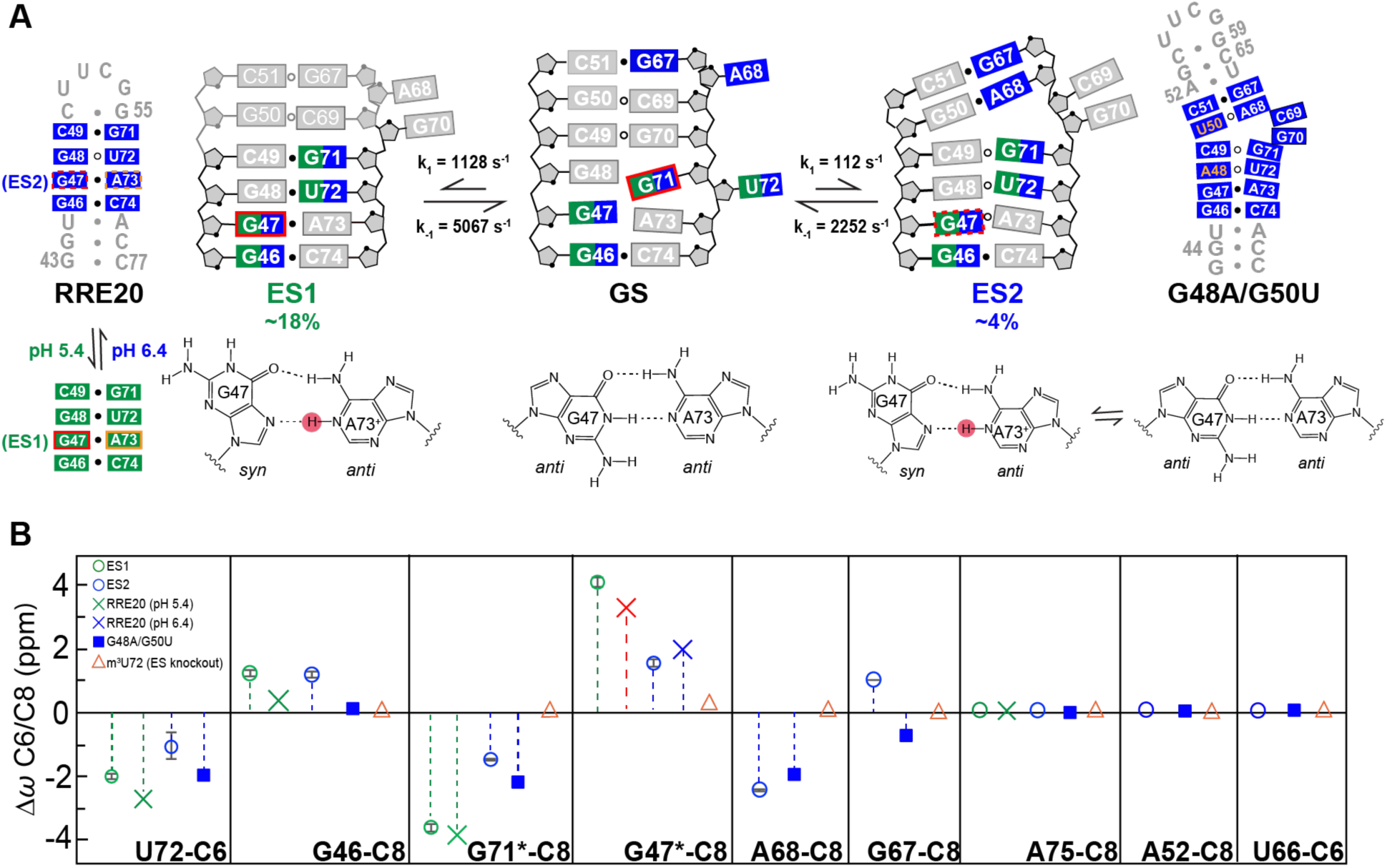
Chemical shift fingerprinting RRE ESs. (A) Putative secondary structure of RREIIB ES1 and ES2 along with ES1 (RRE20) and ES2 (G48A/G50U) stabilizing mutants. Nucleotides that experience exchange due to ES1 and ES2 are colored green and blue, respectively. *syn* bases and protonated adenosine are indicated with red and orange boxes, respectively. The G47(*syn*)-A73^+^(*anti*)⇌G47(*anti*)-A73(*anti*) equilibrium indicated with dashed boxes on ES2. The different G47-A73 conformations in GS, ES1, and ES2 are shown below the secondary structures. Base pairs with and without detectable H-bonds are indicated using filled and empty circles, respectively. (B) Comparison of Δω = ω_ES_ − ω_GS_ obtained using RD (values obtained in the absence of Mg^2+^ indicated using asterisk) with corresponding Δω_mut_ = Δω_ES-mutant_ − ω_wild-type_ obtained for the mutants. Buffer conditions: 15 mM sodium phosphate, 25 mM NaCl, 0.1 mM EDTA, pH 6.4 with or without 3 mM MgCl_2_.

Global fitting of the RD data (see methods) indicates that in the presence of Mg^2+^, ES1 is highly abundant with p_B_ ~18% and exchange rate k_ex_ ~ 6,195 s^−1^ that is fast on the NMR timescale (Fig. S3B, Table S1). Its formation involves changes in G46, G47, G71 and U72 in the lower internal loop (Fig. 2A). Relative to ES1, ES2 has a lower abundance with p_B_ ~ 4% and comparable exchange rate with k_ex_ ~ 2,364 s^−1^. Its formation involves changes in G46 and U72 at the lower internal loop as well as A68 and G67 in and above the internal loop (Fig. 2A). While the similar Δω values indicate that Mg^2+^ does not alter the ES conformation (Fig. S3C), the ES1 population decreased 3-fold in the absence of Mg^2^+ (p_B_= 5.7% and k_ex_ = 10,751 s^−1^) while that of ES2 increased ~3-fold (p_B_ = 11% and k_ex_ = 2,311 s^−1^) (Fig. S3B, Table S1). Thus, Mg^2+^ preferentially stabilizes ES1 over ES2. These Mg^2+^ induced changes in the populations of ES1 and ES2 explain the Mg^2+^-induced changes in line-broadening observed in 2D HSQC spectra (Fig. S2).

### Characterizing ES structures

We analyzed the ES1 and ES2 chemical shifts to gain insight into their conformations (Table S1) (16–18). The chemical shifts provide fingerprints that can be used to assess structural features such as whether or not nucleotides adopt helical versus bulged or *syn* versus *anti* base conformations. Many nucleotides (U72-C6, G71-C8, and G46-C8) had similar chemical shift fingerprints indicating that they adopt similar conformations in ES1 and ES2. For example, U72-C6, Δω_ES1_= −2.3 ppm and Δω_ES2_= −1.2 ppm suggest a bias toward a helical conformation; G71-C8 Δω_ES1_= −3.8 ppm and Δω_ES2_= −16 ppm suggests a change from a *syn* to *anti* base conformation; while for G46-C8 Δω_ES1_ = 1.1 ppm and Δω_ES2_ = 1.1 ppm suggests a less helical conformation (Table S1). The fingerprints for G71-C8 and U72-C6 (Table S1) suggest a more helical conformation in ES1 compared to the GS or ES2.

Other nucleotides have different chemical shift fingerprints indicating that they adopt different conformations in ES1 and ES2. For example, G47-C8 (Δω_ES1_ = 4.0 ppm) suggests a *syn* base conformation in ES1 but a mixture of *syn* and *anti* in ES2 (Δω_ES2_ = 1. 6 ppm) (Table S1). A68-C8 did not show any exchange due to ES1 and Δω_ES2_ = −2.5 ppm suggests a more helical conformation in ES2. No residues showed exchange due to ES1 but not ES2 (Table S1).

MC-fold (81) was used to predict alternative low energy secondary structures for RREIIB. Strikingly, the two lowest energy structures following the GS could explain the ES1 and ES2 chemical shift fingerprints (Fig. S3D). The putative ES1 structure forms by migrating the lower U72 bulge upwards by having U72 displace G71 for pairing with G48 while G71 displaces G70 for pairing with C49, leaving G70 bulged out (Fig. 3A). This putative ES1 structure also features a G48-U72 mismatch and C49-G71 base pair, which can explain the increase in helical character relative to the GS at U72 and G71, respectively, while A68 remains bulged out, explaining lack of ES1 RD at this site. The putative ES1 structure also features a G47-A73 mismatch. If this mismatch were to adopt a Hoogsteen type G(*syn*)-A+(*anti*) conformation, the *syn* base could explain the large downfield-shift at G47-C8 in ES1. That the RD reports on a protonated G(*syn*)-A^+^(*anti*) was verified using pH dependent measurements; increasing the pH to 8 quenched RD at G47-C8 consistent with a protonated G47(*syn*)-A73^+^(*anti*) species (Fig. S4A). Protonation of A73 can also explain broadening of aromatic A73-C8H8 and A73-C2H2 resonances (Fig. S1).

On the other hand, the putative ES2 structure forms by downward migration of the upper stem IIB bulge. Here, A68 displaces C69 for pairing with G50, and this is accompanied by the additional flipping out of G70 possibly to accommodate the junctional G50-A68 (Fig. 3A). This putative ES2 structure can explain the increase in helical character relative to the GS observed at A68 which likely forms G50(*anti*)-A68(*anti*) mismatch based on the chemical shifts. The putative ES2 structure also features a G47-A73 mismatch, which based on the weakly downfield shifted G47-C8 (Δω_ES2_ = 1.6 ppm) is predicted to form a ~50:50 G47(*anti*)-A73(*anti*)⇌μG47(*syn*)-A73^+^(*anti*) dynamic equilibrium. In contrast, to G47-C8, increasing the pH to 8 had a negligible effect on ES2 exchange as judged based on RD measurements on A68-C8 (population changed from ~11% to ~10%, Fig. S4B). This is consistent with the absence of a predominantly protonated mismatch in ES2.

It should be noted that among the predicted secondary structures are close variants of the proposed ES1 and ES2 which feature small variations with respect to which nucleotides are junctional versus unpaired (Fig. S3D). We cannot rule out that such conformations also exist in 10-fold lower abundance relative to the ESs characterized here.

### Testing non-native ES structures using mutate-and-chemical shift fingerprinting

A mutate-and-chemical-shift-fingerprinting strategy (16) was used to test the proposed ES structures (Fig. 3 and S3E). Mutations were introduced to stabilize conformational features unique to the two ESs. The chemical shifts of the mutant relative to wild-type (Δω_mut_) were then compared to those measured by RD (Δω_RD_).

ES1 and ES2 share a similar lower helical stem with G47-A73 and G48-U72 mismatches. This stem was stabilized using a hairpin construct (RRE20) capped by a stable UUCG loop (Fig. 3A). NMR analysis revealed that RRE20 folds into the expected secondary structure (Fig. S5). Very good agreement was observed between Δω_mut_ and Δω_RD_ measured for ES1 and ES2 for U72-C6, G46-C8, and G71-C8 (Fig. 3B and S3E). In addition, sites that showed no detectable RD (A75-C8, A52-C8, U66-C6, and G53-C8) also experienced very small chemical shift perturbations (Fig. 3B and S3E).

ES1 and ES2 differ significantly with regards to the chemical shift fingerprints for G47-C8. At pH 6.4, G47-C8 in RRE20 was only partially downfield shifted (Δω_mut_ = 1.7 ppm) in very good agreement with the ES2 chemical shifts (Δω_RD_ = 1.6 ppm). A strong H8-H1’ NOE cross peak was also observed (Fig. S5) indicating that G47 at least partially adopts the predicted *syn* conformation consistent with the proposed ~50:50 G47(*anti*)-A73(*anti*) ⇌ G47(*syn*)-A73^+^(*anti*) equilibrium. This is notable considering that G-A mismatches can adopt a wide range of pairing geometries (G(*syn*)-A^+^(*anti*), G(*anti*)-A(*syn*), and G(*anti*)-A(*anti*)) depending on sequence, structural context, and pH (82–85). Lowering the pH to 5.4 resulted in the expected fully downfield shifted G47-C8 (3.3 ppm) in good agreement with the RD derived ES1 chemical shifts (Δω_RD_ = 4 ppm) (Fig. 3B and S5). This was also accompanied by a downfield shift in A73-C8 (1.5 ppm) as expected from protonation of adenine-N1 and formation of a predominantly G47(*syn*)-A73^+^(*anti*) conformation in ES1 (Fig. S5). Lowering the pH had little effect on other resonances, indicating that in the mutant RRE20 (but not necessarily the ES), the G47(*anti*)-A73(*anti*) ⇌ G47(*syn*)-A73^+^(*anti*) equilibrium is decoupled from any other conformational changes in the molecule (Fig. S5).

A second mutant (G48A/G50U-RREIIB) was designed to stabilize the non-canonical G48-U72 and G50-A68 mismatches in ES2 (Fig. 3A). NMR analysis indicates that G48A/G50U-RREIIB folds into the putative ES2 secondary structure with G48-U72 and G50(*anti*)-A68(*anti*) mismatches and a dinucleotide C69-G70 bulge (Fig. S6). This provided independent support for proposed dinucleotide bulge in ES2 for which no RD data could be measured. Again, excellent agreement was observed between Δω_mut_ and Δω_RD_ for U72-C6, G46-C8, G71-C8 and A68-C8, and (Fig. 3B and S3E). The only exception was G47-C8 for which the chemical shifts and NOEs indicate that the G47-A73 adopts an *anti-anti* conformation with chemical shifts similar to the GS. Thus, the G48A/G50U-RREIIB mutant does not appear to capture the ES2 of G47-A73. This is not surprising considering that the neighboring residue is mutated and *syn:anti* equilibria are known to be quite dependent on sequence context (86).

Finally, we also tested a mutant designed to destabilize both ES1 and ES2 through introduction of the chemically modified m^3^U72 (m^3^U72-RREIIB) that blocks formation of the G48-U72 mismatch in ES1 and ES2 without affecting the GS in which U72 is bulged out (Fig. S7A). As expected, 2D NMR spectra of m^3^U72-RREIIB were very similar to their unmodified counterparts, with minor differences observed in and around the site of modification (Fig. 3B and S3E). Interestingly, some resonances such as G71 shifted toward the GS (Fig. S7A). The modification also reduced line broadening in 2D NMR spectra at G48, G71 and A73 exactly as expected from destabilization of both ES1 and ES2 (Fig. S7B). This provides independent support that U72 is paired in ES1 and ES2.

### Characterizing ESs in the RREII three-way junction

Examining whether or not the ESs also form in the large RREII construct (68 nt) is complicated by severe spectral overlap and low sensitivity in this larger RNA particularly when resonances are broadened due to conformational exchange. Recognizing that the ES1 population in RREIIB is quite high (p_B_ ~18%), we reasoned that it may be possible to directly observe ES1 in RREII provided that specific sites were ^13^C/^15^N labeled to aid observation of resonances belonging to the minor ES populations from the overwhelming excess of resonances belonging to the dominant GS. If exchange in RREII was slow on the NMR timescale, it should be feasible to directly observe the chemical shifts of ES1 and thereby also estimate its population (Fig. 4). If on the other hand exchange was fast on the NMR timescale, as is observed in RREIIB (Δω/k_ex_ = 2.4), the population weighted chemical shifts for nucleotides that experience large Δω would be shifted either toward the GS or ES, in a manner dependent on their relative populations. The strategy therefore affords the possibility to detect ESs in large RNAs under both fast and slow exchange conditions.

**Figure 4.**
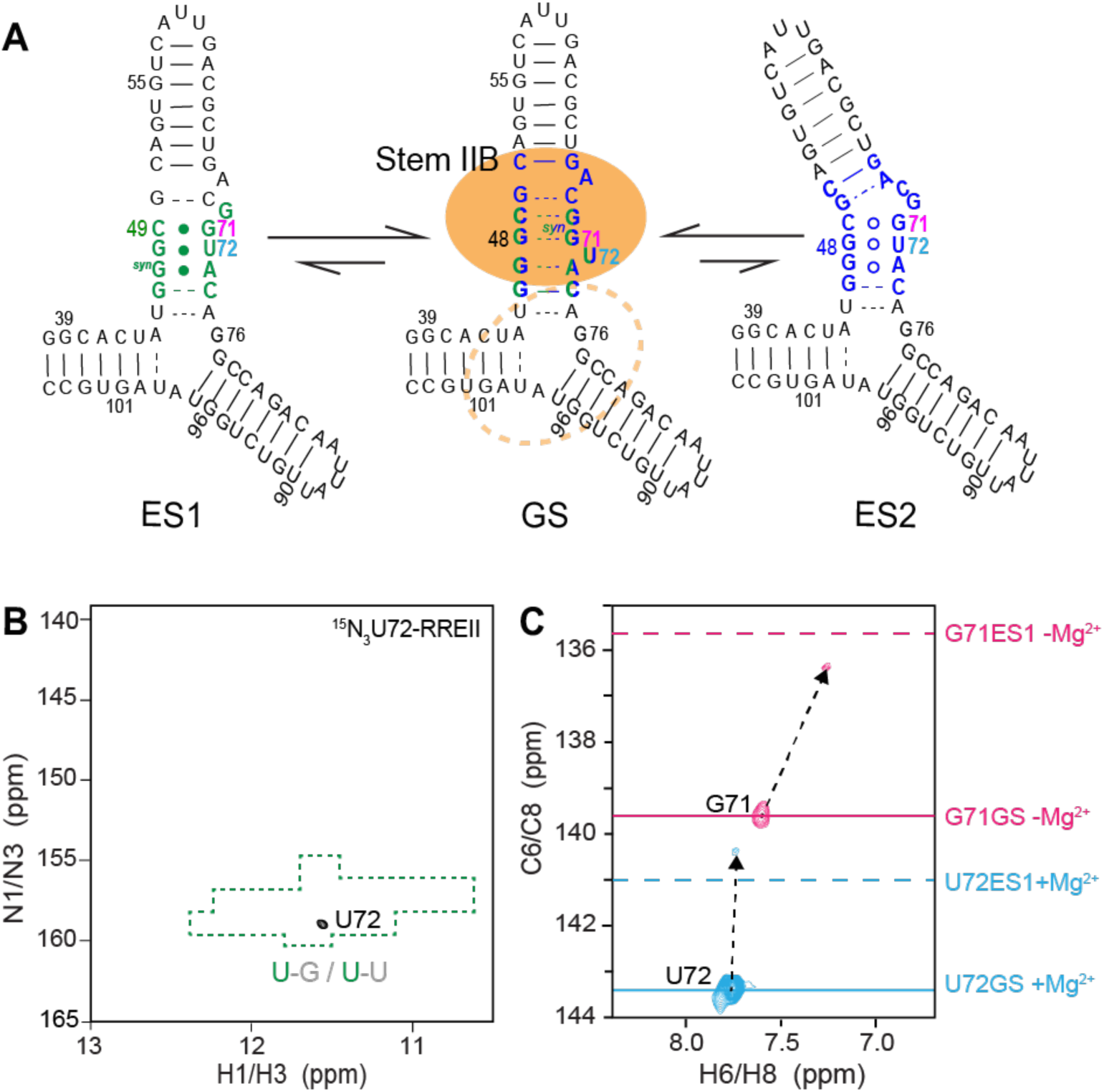
Selective labeling strategy to probe conformational exchange in RREII. (A) Proposed secondary structure for ES1 and ES2 in RREII based on the ESs observed in RREIIB. Nucleotides that experience exchange due to ES1 and ES2 are colored green and blue, respectively. Rev primary and secondary binding sites are highlighted using a filled and dashed orange circle, respectively. (B) 2D NH HSQC spectrum of site-specifically labeled RREII at 25 °C in 3 mM Mg^2+^ showing a single imino resonance at the characteristic chemical shift region (highlighted in green) expected for a G-U wobble. (C) 2D CH HSQC spectra of site-specifically labeled RREII at 25 °C showing the appearance of major resonances consistent with the GS in RREIIB (solid line) and minor resonances consistent with ES1 in RREIIB (dashed line). Spectra for G71 was measured in the absence of Mg^2+^. Buffer conditions: 15 mM sodium phosphate, 25 mM NaCl, 0.1 mM EDTA, pH 6.4 with or without 3 mM MgCl_2_.

The U72 imino resonance is not observable in the GS because it adopts an unpaired conformation (Fig. 4A). In contrast, not only should this resonance be observable in both ES1 and ES2, it is predicted to resonate at specific NMR chemical shifts unique to the G-U wobble (Fig. 4A). We recorded NMR spectra for ^15^N3-U72-RREII in the presence of 3 mM Mg^2+^. Under these conditions, and based on RD measured in RREIIB, we expect ES1 p_B_ ~18% and ES2 p_B_ ~ 4%. Strikingly, NMR spectra of ^15^N3-U72-RREII revealed a single uridine imino resonance with ^1^H and ^15^N chemical shifts that are characteristic of a G-U wobble (Fig. 4B). To further confirm direct observation of ES1 in RREII, we selectively labeled ^13^C6-U72 and observed a major resonance corresponding to the GS, in which U72 is bulged out, and a minor upfield-shifted resonance (−2.7 ppm) that is in excellent agreement with the chemical shift measured for ES1 using RD (Δω_RD_ = −2.3 ± 0.1 ppm) and in poorer agreement with ES2 (Δω_RD_ = 1.2 ± 0.5 ppm) (Fig. 4C). Finally, we also prepared a sample with labeled ^13^C8-G71 and observed the expected major resonance corresponding to the GS in which G71 forms G48-G71(*syn*) and a minor upfield-shifted (−4.1 ppm) resonance which presents an *anti-base* of G71 in excellent agreement with ES1 chemical shift (Δω_RD_ = −3.8 ± 0.2 ppm) and poorer agreement with ES2 (Δω_RD_ = −1.6 ± 0.05 ppm) (Fig. 4C). These spectra were recorded in the absence of Mg^2+^ to allow comparison with RD data on RREIIB and under these conditions we expect ES1 p_B_ ~ 6% and ES2 p_B_ ~ 11%.

The population of ES1 estimated from the integrated volumes of the resonances is ~20% in absence or presence of 3 mM Mg^2+^. This is in good agreement with ES1 population (~18%) measured in RREIIB in the presence of 3 mM Mg^2+^. Taken together, these data show that ES1 also exists in RREII three-way junction with similar populations as RREIIB and that the three-way junction decouples the dependence of the ES1 population on Mg^2+^. While we do not observe ES2 resonances, we cannot rule that the exchange is too fast or that the ES2 population falls below detection limit in this experiment. These results establish site-specific labeling as a new methodology for directly observing ESs in large RNAs.

### Rev peptide binds the RRE ESs with significantly lower affinity

The ES1 and ES2 conformations disrupt structural elements in stem IIB that have previously been shown to be critical for Rev binding. The hypothesis that Rev ARM peptide binds more weakly to the non-native ES1 and ES2 was tested using a mutate- and-rescue strategy. The Rev-ARM/stem IIB interaction has been shown to recapitulate the affinity and specificity of the full complex (48,87). We tested the impact of mutations that stabilize ES1 and/or ES2 on binding of fluorescein labeled Rev-ARM peptide (Rev-Fl) using a fluorescence polarization (FP) assay (78). The ES-stabilizing mutants used in NMR chemical shift fingerprinting eliminate structural elements required for Rev binding and also do not afford the possibility to introduce rescue mutations. We therefore designed a new set of ES-stabilizing mutants that preserve key elements required for Rev binding and that could be rescued with compensatory mutations (Fig. 5A). This was not a simple task given the need to avoid mutating residues important for Rev recognition.

**Figure 5.**
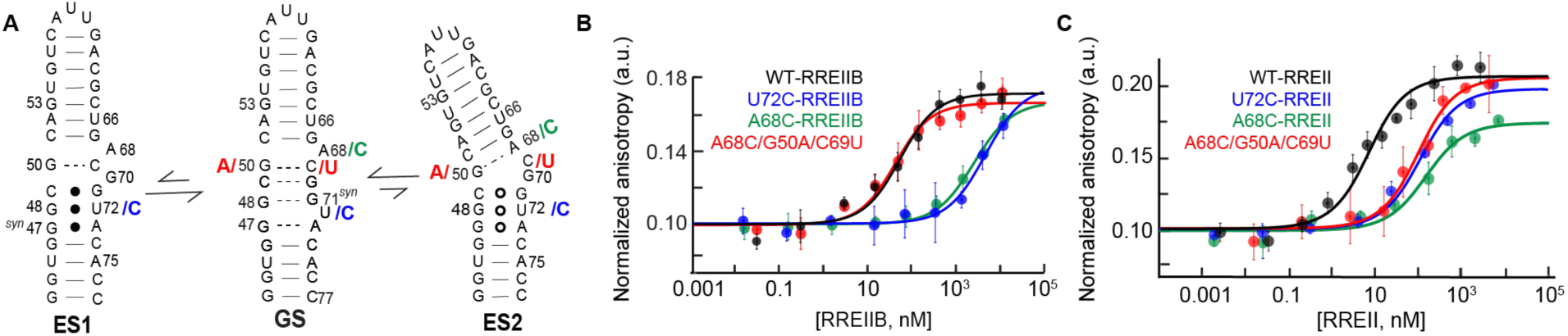
Measurement of the binding affinity between Rev-ARM peptide and RRE, ES-stabilizing mutants, and rescue mutants using fluorescence polarization. (A) The designed ES-stabilizing, ES2-stabilizing and ES2-rescue mutants are colored in blue, green, and red, respectively. (B) Normalized anisotropy values measured for RREIIB and RREII fitted with one-site binding model (see methods). The anisotropy value observed in the absence of RNA was normalized to 0.1. Uncertainty reflects the standard deviation from three independent measurements. Buffer conditions: 30 mM HEPES, pH= 7.0, 100 mM KCl, 10mM sodium phosphate, 10 mM ammonium acetate, 10 mM guanidinium chloride, 2 mM MgCl_2_, 20 mM NaCl, 0.5 mM EDTA, and 0.001% (v/v) Triton-X100. Concentration of Rev-Fl peptide was 10 nM and 1 nM for RREIIB and RREII respectively.

ES2 was stabilized using the A68C point substitution mutation, which replaces the non-canonical G50-A68 mismatch with a canonical G50-C68 Watson-Crick base pair. Both ES1 and ES2 were stabilized using the U72C point substitution mutation, which replaces the G48-U72 wobble with a canonical G48-C72 Watson-Crick base pair (Fig. 5A). While these mutations were predicted to fold into the ESs as the energetically preferred conformation, the NMR spectra show significant line-broadening specifically at nucleotides (U72 and A68) involved in the conformational exchange (Fig. S8A and S8B). These data indicate that the mutants push the conformational equilibrium toward the ESs (to ~50% population) but to a smaller degree than the more stringent mutations used in mutate-and-chemical-shift-fingerprinting (Fig. 3A). We also cannot rule out that the mutations populate other higher energy ESs (Fig. S3D) that may contribute to the observed line-broadening.

As a positive control, Rev-Fl peptide binds to wild-type RREIIB with K_d_ = 45.9 ± 15.2 nM (Fig. 5B and Table 1) in good agreement with prior studies (~30–50 nM) (48,87). The binding affinity (K_d_ > 2,000 nM) decreased 40 to 80-fold for the U72C and A68C RREIIB mutants (Table 1). Because of the reduced binding affinity, binding did not reach saturation, and increasing the concentration of the ES-stabilized mutants further resulted in additional low-affinity binding (88) (Fig. S9A). To test whether the reduction in binding affinity is due to stabilization of the unique ES conformation, we introduced additional rescue mutations (A68C/G50A/C69U-RREIIB; Fig. 5A) designed to restore the GS conformation by stabilizing an A50-U69 canonical base pair in the mutant. Indeed, the rescue mutant restored Rev binding to wild-type levels (K_d_ = 33.0 ± 11.3 nM) (Fig. 5B and Table 1). Thus, the lower binding affinity observed for the ES-stabilizing mutants is due to their alternative conformations and not due to changes in sequence.

**Table 1.**
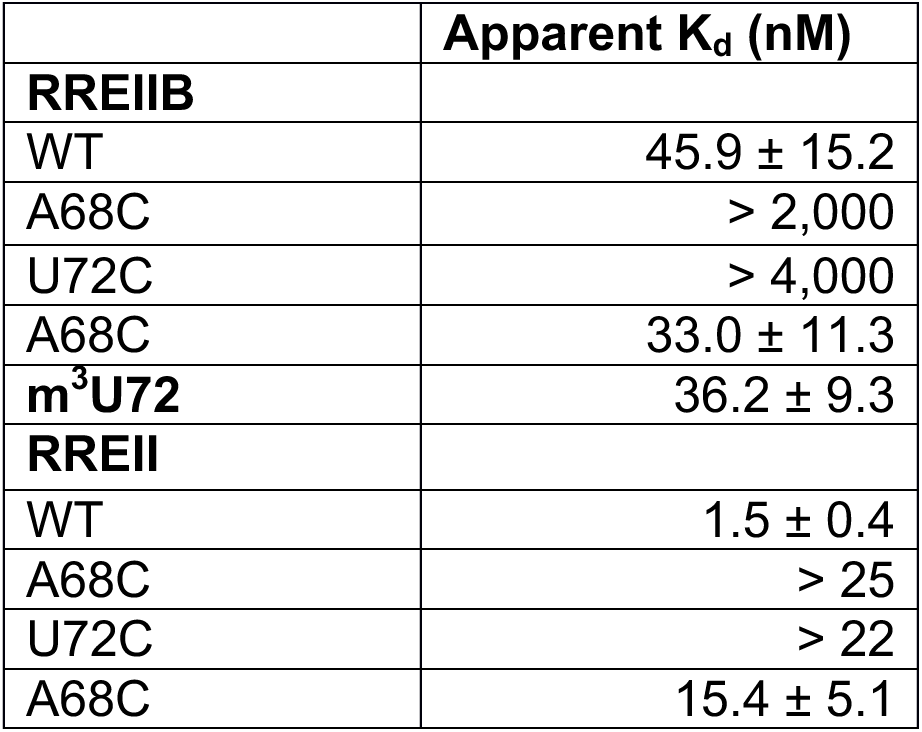
The apparent K_d_ describing binding of wild-type RREIIB and RREII and their mutants to Rev-Fl obtained by fitting binding curves to a one-site binding model. Uncertainty represents standard deviation over triplicate measurements. The concentration of Rev-Fl peptide is 10 nM and 1 nM for RREIIB and RREII, respectively. Buffer conditions: 30 mM HEPES, pH= 7.0, 100 mM KCl, 10mM sodium phosphate, 10 mM ammonium acetate, 10 mM guanidinium chloride, 2 mM MgCl_2_, 20 mM NaCl, 0.5 mM EDTA, and 0.001% (v/v) Triton-X100.

| | Apparent $K_d$ (nM) |
| --- | --- |
| **RREIIB** |  |
| WT | $45.9 \pm 15.2$ |
| A68C | $> 2,000$ |
| U72C | $> 4,000$ |
| A68C | $33.0 \pm 11.3$ |
| **m<sup>3</sup>U72** | $36.2 \pm 9.3$ |
| **RREII** |  |
| WT | $1.5 \pm 0.4$ |
| A68C | $> 25$ |
| U72C | $> 22$ |
| A68C | $15.4 \pm 5.1$ |

The ES-stabilizing mutations also weakened binding to the three-way junction RREII (Fig. 5C). As a positive control, Rev-Fl binds to wild-type RREII with 30-fold higher affinity (K_d_ = 1.5 ± 0.4 nM) compared to RREIIB, consistent with prior studies which report affinities on the order of ~3–9 nM (Table 1) (41,75). Binding to the A68C and U72C RREII ES stabilizing mutants was reduced ~15 fold (K_d_ > 20 nM, Table 1). Once again, because of the low binding affinity, the binding curves did not reach saturation and further addition of ES-stabilizing mutants resulted in additional low-affinity binding (Fig. S9A). In RREII, the A68C ES2 rescue mutant (A68C/G50A/C69U-RREII) only partially restored wild-type binding (K_d_ = 15.5 ± 5.1 nM). The binding affinity measured for the rescue mutant was more similar to that observed for the same rescue mutant in RREIIB (K_d_ = 33.0 ± 11.3 nM) as compared to the wild-type sequences where the differences were 30–fold. This indicates that the higher affinity to RREII relative to RREIIB may be linked to nucleotides G50 and C69 used in the rescue mutations.

Finally, we measured the binding to ES-destabilizing mutant m^3^U72-RREIIB. Increasing the binding-competent GS from 80% to 100% is predicted to improve the binding and decrease the observed K_d_ by a factor of 1.25. Strikingly, the binding affinity (K_d_ = 36.2 ± 9.3 nM, Fig. S9B and Table 1) did indeed increase by ~1.25 fold as predicted in the ES knockout mutant as compared to wild-type RREIIB.

## DISCUSSION

Our study adds to a growing view that non-canonical regions of RNA do not fold into a single secondary structure but rather exist as a dynamic equilibrium of alternative conformations that have non-native secondary structure (16–19). Relative to other RNA ESs, which tend to have populations on the order of ~1%, the ES population in RREII is ~ 20% in the presence of Mg^2+^. Given that the RRE ESs bind Rev peptides with 15 to 80-fold weaker affinity, these results underscore how iso-energetic structures of RNA can have very different biological activities.

The high population of the ESs also implies that they could contribute to chemical probing data, which is averaged over the ensemble. In this regard, the RRE ensemble helps clarify the high SHAPE reactivity reported in previous studies for many nucleotides within the internal loop of stem IIB, which are base paired in the GS (42,89). For example, in ES1, G70 flips out and G47 adopts a non-canonical *syn* conformation, and this could explain the high reactivity at these nucleotides (Fig. 6A) (42,89). Importantly, while the ES ensemble predicts that many nucleotides will be unpaired at a given point in time, they are seldom unpaired simultaneously. This emphasizes the importance of interpreting chemical probing data in terms of dynamic ensembles (30,31). Ensemble-based approaches (29,30,90) for interpreting chemical probing data may help clarify the nature of the stem II RRE ensemble within the complex cellular environment. Indeed, the ensemble cannot explain the high SHAPE reactivity at G46-C74 (89), which is paired in all conformations. Thus, we cannot rule out that the RRE ensemble differs in the complex cellular environment as compared to *in vitro*.

**Figure 6.**
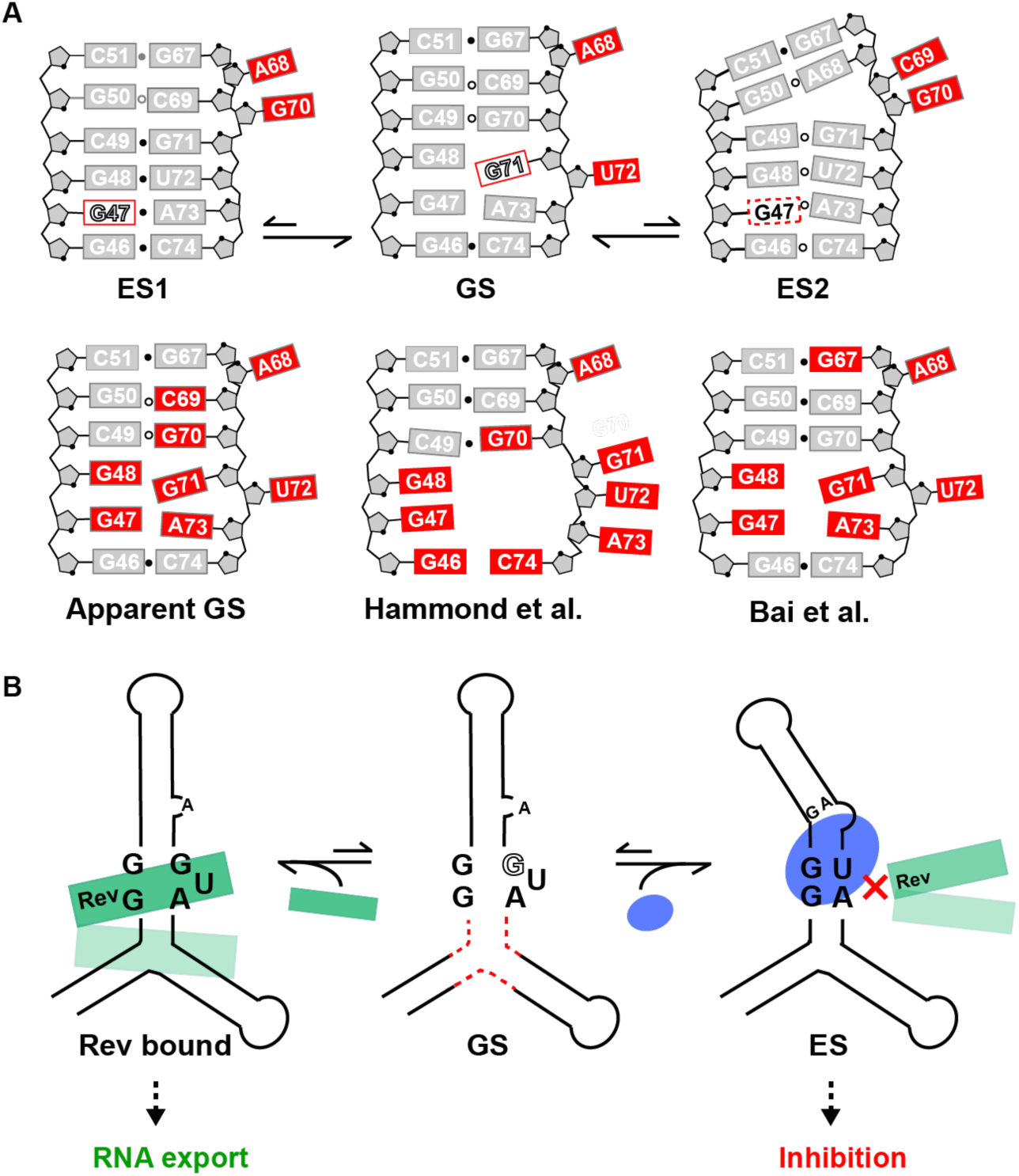
Implications of RRE dynamic ensemble. (A) The RRE ES ensemble results in different conformations that expose different nucleotides for reactivity. This could contribute to high reactivity (apparent GS) at several nucleotides, including ones that may form canonical pairs in the GS. Results are compared with SHAPE reactivity measured previously by Hammond et al. (89) and Bai et al. (42). The *syn* base in GS and ES1 are indicated using red rectangles and open letters. Residues in *syn* to *anti* conformational exchange are indicated using dashed rectangles. (B) Proposed role for RRE ESs in providing mechanisms for conformational cooperativity during Rev (in green) recognition by organizing the secondary binding site (dashed red lines). Stabilization of ESs using small molecules (in blue) could provide a strategy for inhibiting Rev binding and viral RNA export for anti-HIV therapeutics. The *syn* base is indicated using open letters.

The addition of the three-way junction did induce small perturbations at stem IIB (Fig S8A and S8B). In addition, mutations to stem IIB that stabilize the ESs induced changes in chemical shifts outside stem IIB in the RREII three-way junction (Fig. S8C). Thus, it is possible that formation of the ESs in stem IIB, are correlated with other conformational changes at the three-way junction region. Because of this conformational coupling, binding of Rev to stem IIB could help organize the second binding site at the three-way junction, including possibly by diminishing the population of any ESs, and thereby provide an additional source of binding cooperativity based on RNA conformational dynamics. Indeed, prior studies showed that replacement of the three-way junction with a stable duplex leads to loss of binding cooperativity (48). Further studies are needed to test this hypothesis. In this regard, the new site-specific labeling strategy introduced in this work provides a new means by which to extend our studies and examine ESs in the RRE stem II three-way junction.

The RRE ensemble also helps with the interpretation of the extensive mutagenesis studies done on the RRE stem IIB (58,60,91,92). These mutations have been shown to inhibit Rev binding by 60 to 2000-fold (58,60,91,92) and to suppress activity in cell based assays by up to 90% (58,60). Interestingly, none of these prior mutations are predicted to stabilize ES1 or ES2. Rather, they are predicted to disrupt key nucleotides (G47, G48, C49, G70, G71, A73) required for Rev binding. Thus, the reduced binding affinity or activity observed with these prior mutations most likely reflects changes in the contacts and/or conformation of the RRE GS, and not stabilization of alternative ESs.

The RRE stem IIB has been the subject of much effort directed toward the development of small molecule anti-HIV therapeutics (36–38,40). Most of these studies have relied on high throughput screening and/or the rational design of inhibitors that target the RRE GS (36–38,40). The new non-native ESs, which remodel key structural features required for Rev binding could constitute novel states for targeting with small molecules. Determining the structures of the RRE ESs would make it possible to apply ensemble-based virtual screening approaches (12,13) to identify small molecule that selectively bind the ESs. The ES-stabilizing mutants could also be subjected to high throughput screening to help identify compounds that selectively bind these non-native states. Because of their unique allosteric mode of action, the hits emerging from such studies might offer complimentary path toward development of anti-HIV therapeutics relative to approaches targeting the dominant GS.

## Supporting information

## SUPPLEMENTARY DATA

Supplementary Data are available at NMR online.

## ACKNOWLEDGEMENT

We thank the Duke Magnetic Resonance Spectroscopy Center for supporting NMR experiments, and thank Duke Center of RNA Biology for supporting fluorescence polarization measurement.

## FUNDING

This work is supported by US National Institutes of Health [P50 GM103297 to H.M.A.] and the Austrian Science Fund [P28725 and P30370 to C.K.]. Funding for open access charge: US National Institutes of Health [P50 GM103297].

## CONFLICT OF INTEREST

H. M.A. is an advisor to and holds an ownership interest in Nymirum Inc., an RNA-based drug discovery company.

