## Supplementary material for "Dynamic ensemble of HIV-1 RRE stem IIB reveals non-native conformations that disrupt the Rev binding site"

### Supplementary data

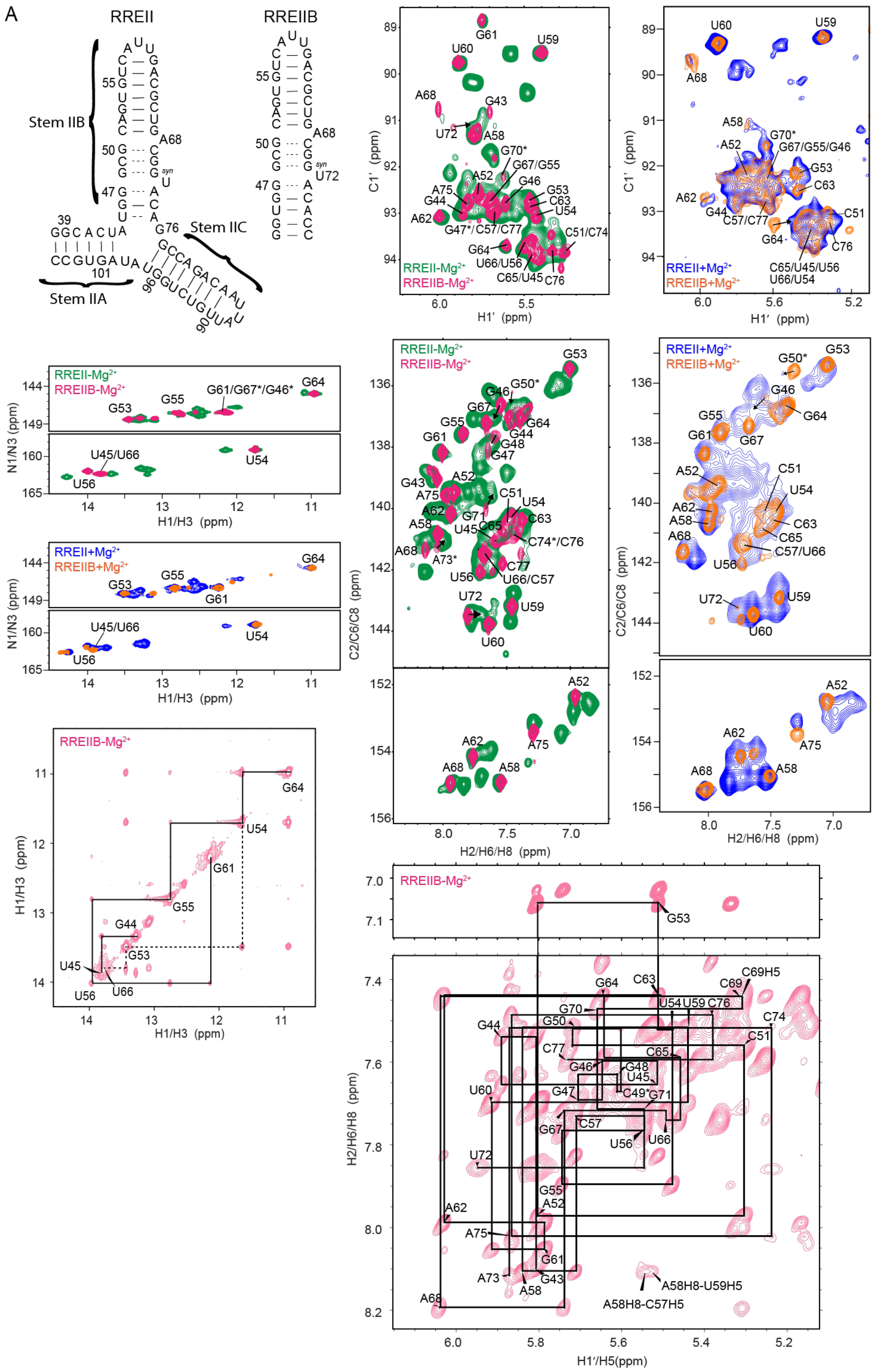

A
RREII
RREIIB
Stem IIB
Stem IIC
Stem IIA
RREII-Mg2+
RREIIB-Mg2+
RREII+Mg2+
RREIIB+Mg2+
C1' (ppm)
H1' (ppm)
N1/N3 (ppm)
H1/H3 (ppm)
C2/C6/C8 (ppm)
H2/H6/H8 (ppm)
H1'/H5(ppm)
A58H8-U59H5
A58H8-C57H5

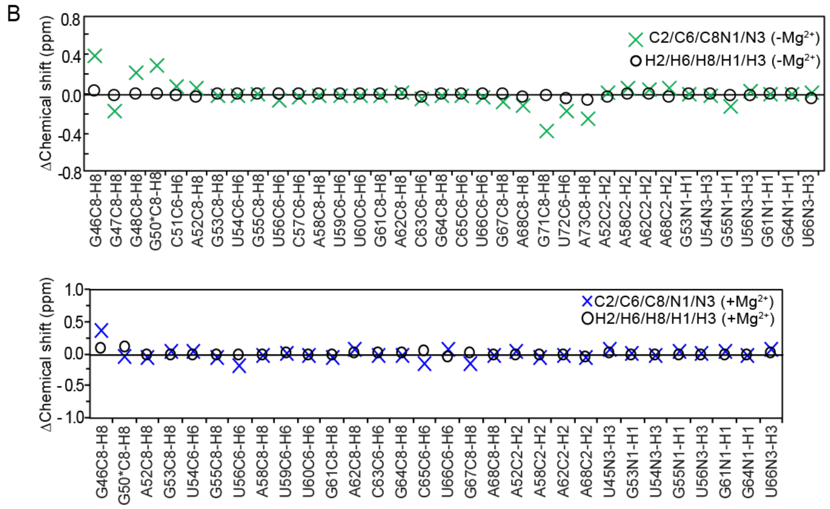


**Figure S1.** NMR spectra measured for RREIIB and RREII. (A) 2D $^1H$-$^1H$ NOESY spectra of exchangeable (10 ºC in 10% $D_2O$ and mixing time 180 ms) and nonexchangeable (37 ºC in 100% $D_2O$ and mixing time 220 ms) protons showing distance based connectivity highlighted on the secondary structure. Base pairs with and without detectable H-bonding are indicated using solid and dashed lines, respectively. Also shown are overlays of 2D NH and CH HSQC spectra for RREIIB and RREII recorded at 25 (-$Mg^{2+}$) and 10 ºC (+$Mg^{2+}$) respectively. Resonances with potentially ambiguous resonance assignment are labeled with an asterisk. (B) Differences between RREIIB and RREII chemical shifts measured in the absence and presence of $Mg^{2+}$. Nucleotides with resonances showing large perturbations are highlighted using red circles on the secondary structure. Buffer conditions: 15 mM sodium phosphate, 25 mM NaCl, 0.1 mM EDTA, pH 6.4 with or without 3 mM $MgCl_2$.

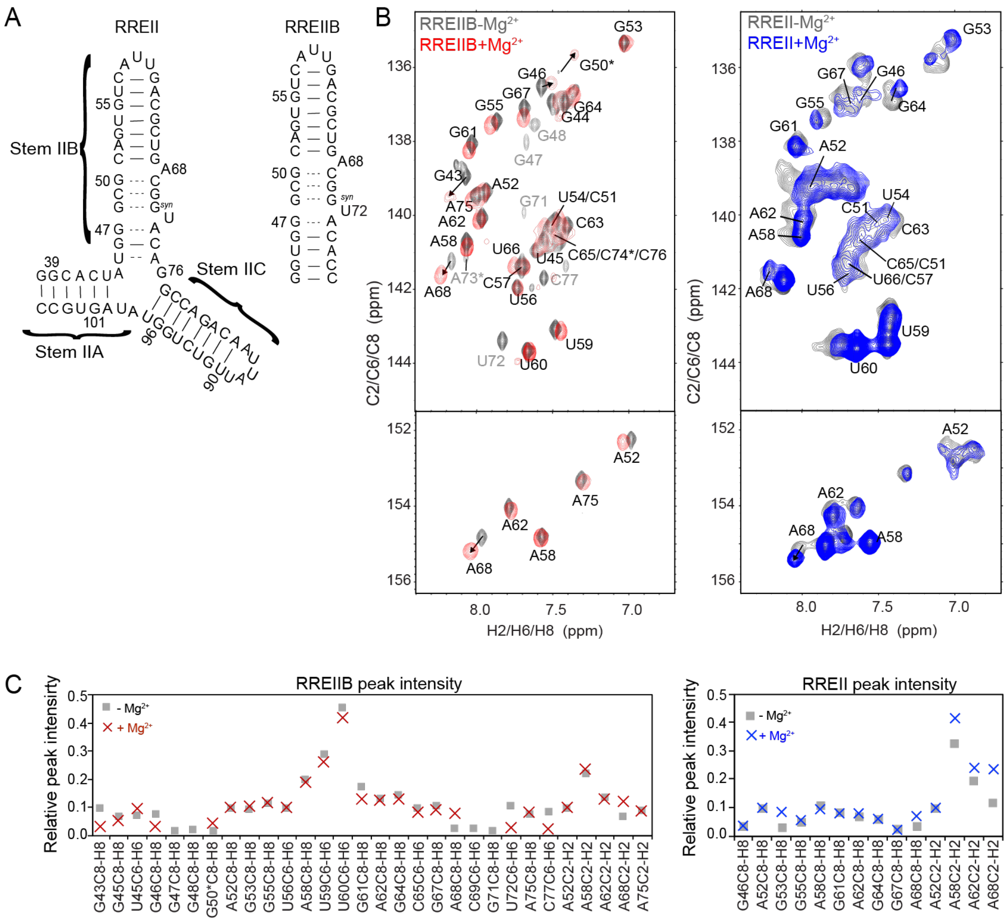


**Figure S2.** Impact of $Mg^{2+}$ on spectra of RREIIB and RREII. (A) Secondary structure of RREIIB and RREII. (B) Comparison of 2D CH HSQC spectra measured for RREIIB (25 ºC) and RREII (10 ºC). (C) Normalized resonance intensities measured for RREIIB (at 25 ºC) and RREII (at 10 ºC) in the presence and absence of 3 mM $Mg^{2+}$. A52-C8H8, A52-C2H2 and U56-C6H6 were used as a reference and normalized to 0.1. Buffer conditions: 15 mM sodium phosphate, 25 mM NaCl, 0.1 mM EDTA, pH 6.4 with or without 3 mM $MgCl_2$.

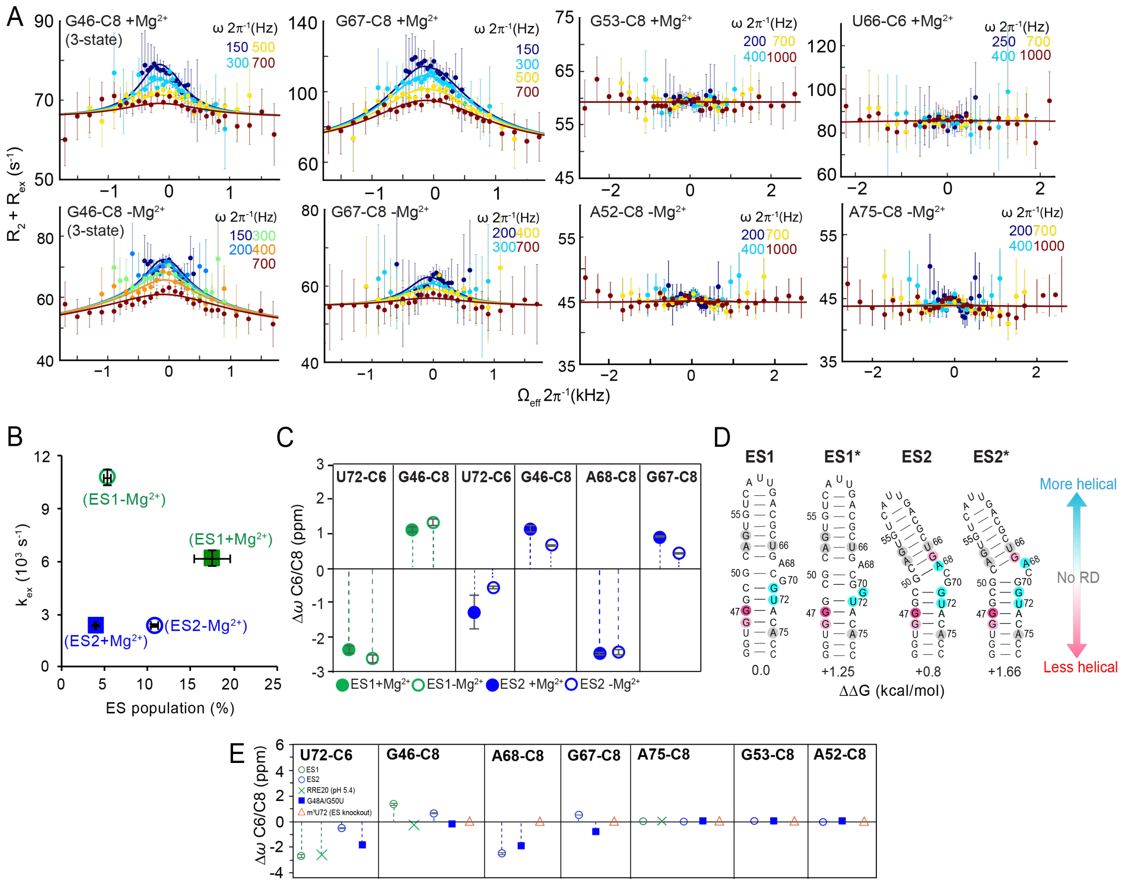

**Figure S3.** RD profiles and exchange parameters measured at 25 ºC. (A) RD profiles measured for RREIIB in the absence and presence of $Mg^{2+}$. (B) ES population and exchange rate measured for RREIIB ES1 and ES2 in the presence (filled) and absence (open) of $Mg^{2+}$. (C) As in main Fig. 3B showing $\Delta\omega_{RD}$ in the absence and presence of $Mg^{2+}$. (D) Chemical shift fingerprints annotated on the putative structural models for ES1 and ES2 obtained as the top two (following the GS) structures predicted by MC-fold (1). The residues on RREIIB with RD measurements are indicated in circles. Gray: no dispersion, cyan: $\Delta\omega < 0$, red: $\Delta\omega > 0$. (E) Comparison of $\Delta\omega$ obtained for the mutants and using RD in the absence of $Mg^{2+}$. Buffer conditions: 15 mM sodium phosphate, 25 mM NaCl, 0.1 mM EDTA, pH 6.4 with or without 3 mM $MgCl_2$.

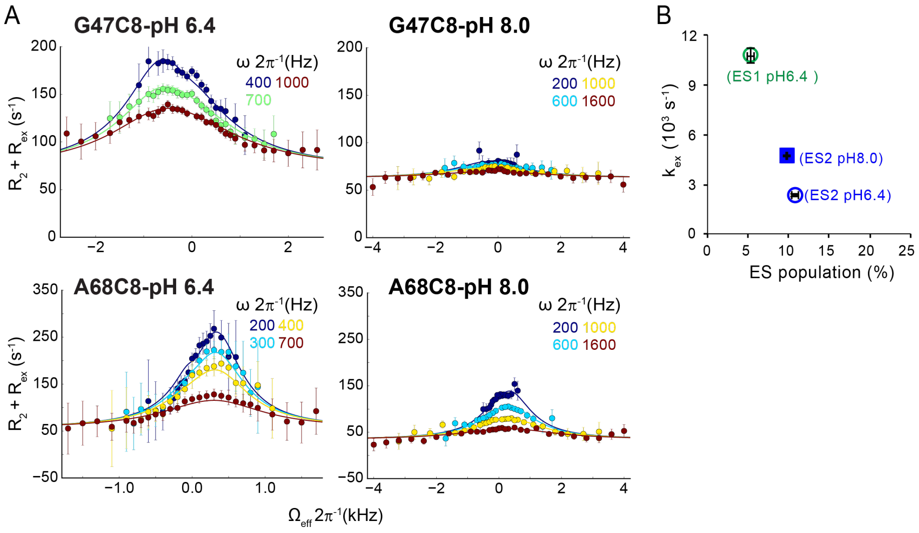


**Figure S4.** pH dependence of $R_{1\rho}$ RD profiles at 25 ºC. (A) $R_{1\rho}$ RD profiles acquired with different color-coded power levels. Solid lines represent the global fits to the RD data (G47-C8 and A68-C8 at pH 6.4 and pH 8.0). Error bars represent experimental uncertainty based on Monte Carlo analysis of monoexponential decay curves and the signal noise. (B) Exchange parameters for ES1 from 3-state fit of $R_{1\rho}$ RD profiles for G47-C8. Note that ES1 is undetectable at high pH and $p_B$ is estimated to be <0.01%, Buffer conditions: 15 mM sodium phosphate, 25 mM NaCl, 0.1 mM EDTA, pH 6.4.

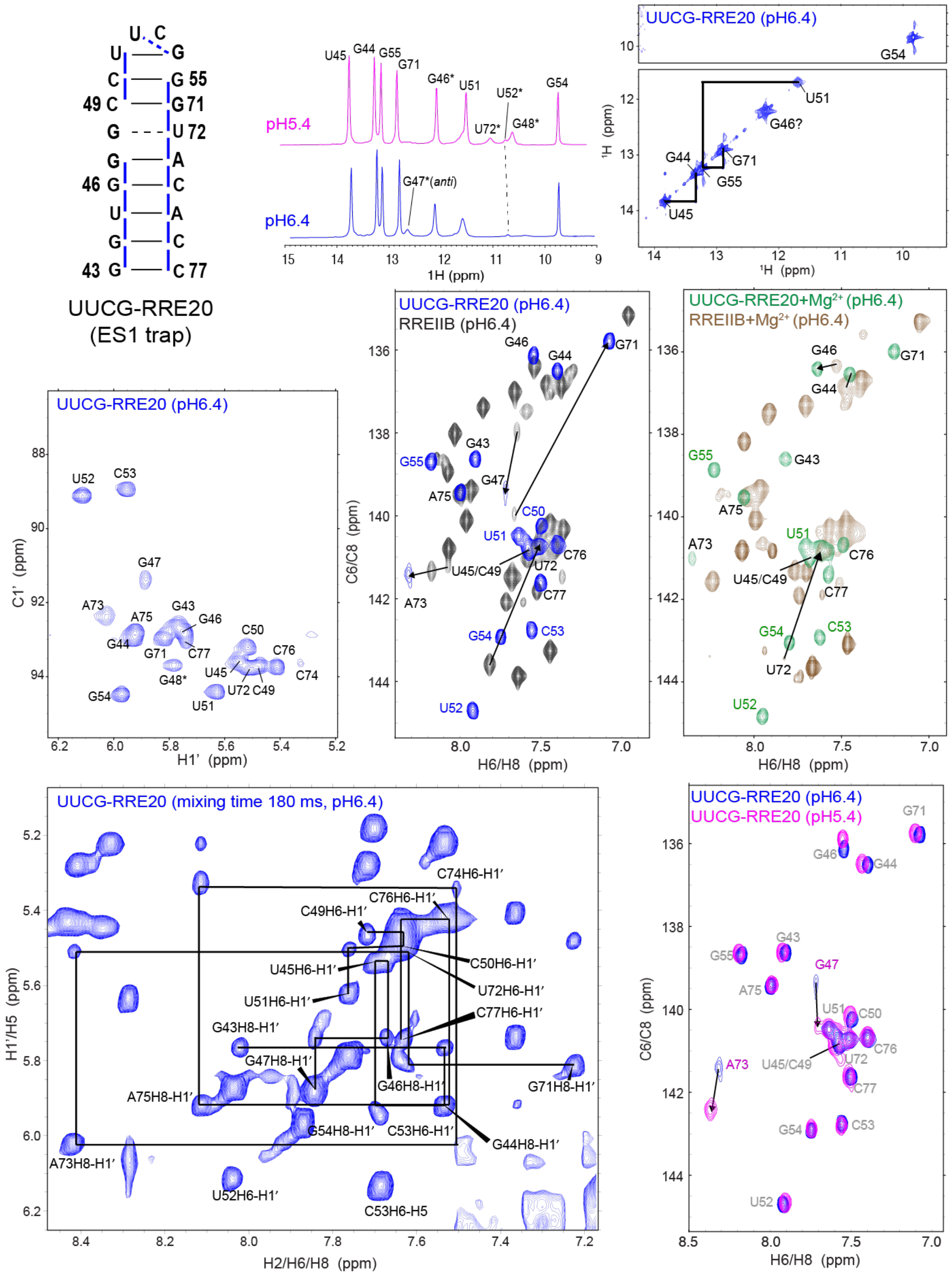


**Figure S5.** NMR spectra and assignments for ES1 stabilizing mutant UUCG-RRE20. Secondary structure of UUCG-RRE20 based on NMR data at 25 ºC. At pH 6.4, a weak G47-H1 imino resonance is observed consistent with an *anti* conformation that exchanges with a *syn* conformation (in which imino is not detectable due to solvent exchange). This resonance was broadened out of detection at pH 5.4 consistent with stabilization of a dominant G47(*syn*) conformation. The imino resonances for the adjacent G48-U72 wobble were not observed at pH 6.4, likely due to conformational exchange in the adjacent G-A mismatch, but were observable at pH 5.4 in which the exchange at the adjacent G-A mismatch is reduced. Uninterrupted intra- and inter-nucleotide H8/6-H1' NOEs were observed for all other resonances except G48 (which is broadened out of detection due to exchange) indicating that these nucleotides adopt a helical conformation. Resonances with potentially ambiguous resonance assignment are labeled with an asterisk. Buffer conditions: 15 mM sodium phosphate, 25 mM NaCl, 0.1 mM EDTA, with or without 3 mM $MgCl_2$.

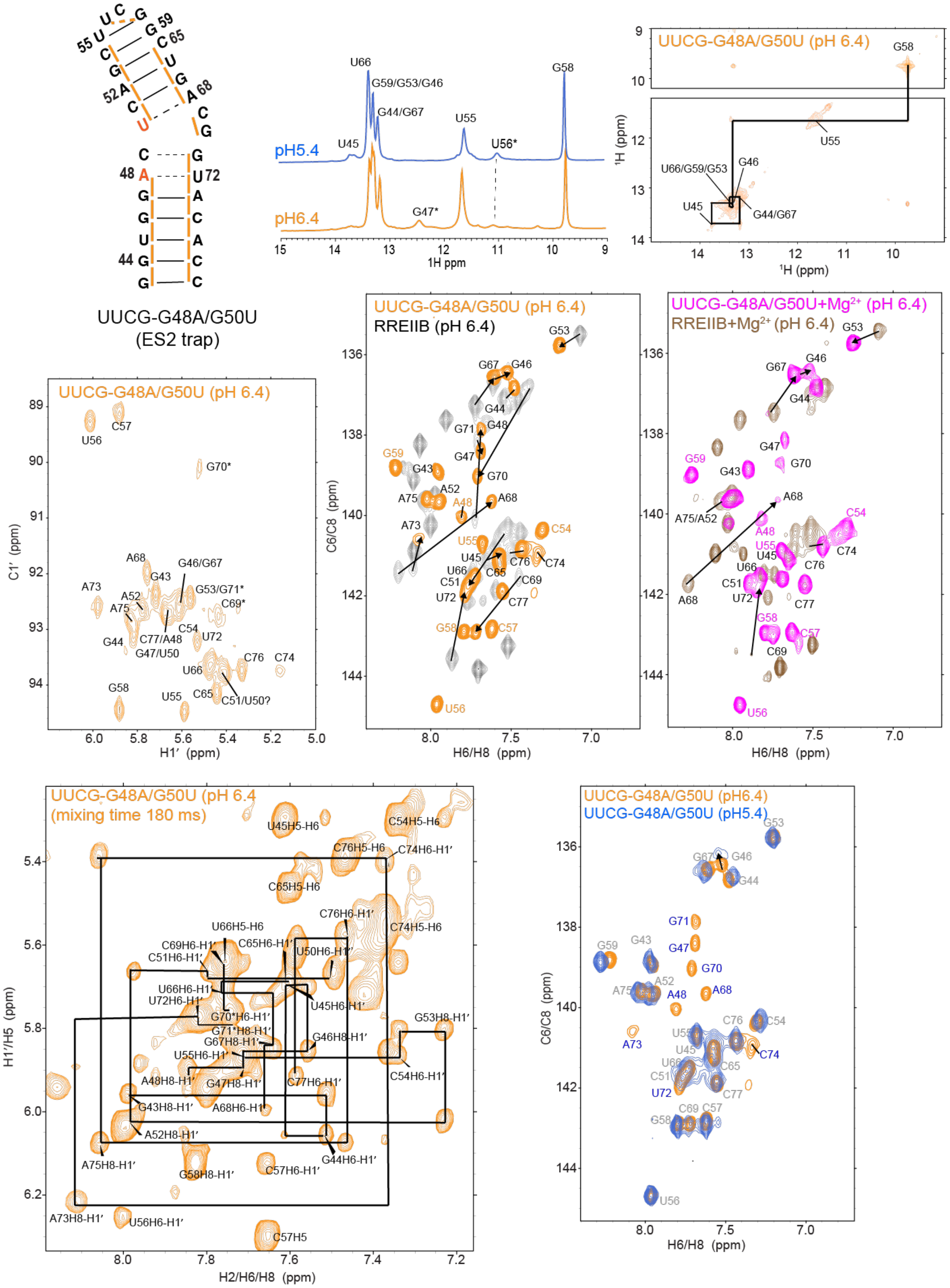


**Figure S6.** NMR spectra and assignments for ES2 stabilizing mutant (UUCG-G48A/G50U) at 25 ºC. Secondary structure of UUCG-G48A/G50U based on NMR data. Base pairs with and without detectable H-bonds are indicated using solid and dashed lines, respectively. The imino and amino resonances for the base pairs around the C69 and G70 bulge were not observed likely due to increase solvent exchange and possibly also conformational exchange. The G47-H1 imino resonance G47H8-H1' NOEs consistent with a G47(*anti*)-A73*(anti*) base pair were observed at pH 6.4 while the imino resonance was broadened significantly at pH 5.4, consistent with the behavior seen with the UUCG-RRE20 mutant. For other non-exchangeable proton, the NOE connectivity breaks at A68-C69 and G70-G71 consistent with C69 and G70 adopting a bulge conformation, whereas uninterrupted intra- and inter-nucleotide H8/6-H1' NOEs were observed for all other nucleotides consistent with the proposed secondary structure. Buffer conditions: 15 mM sodium phosphate, 25 mM NaCl, 0.1 mM EDTA, with or without 3 mM $MgCl_2$.

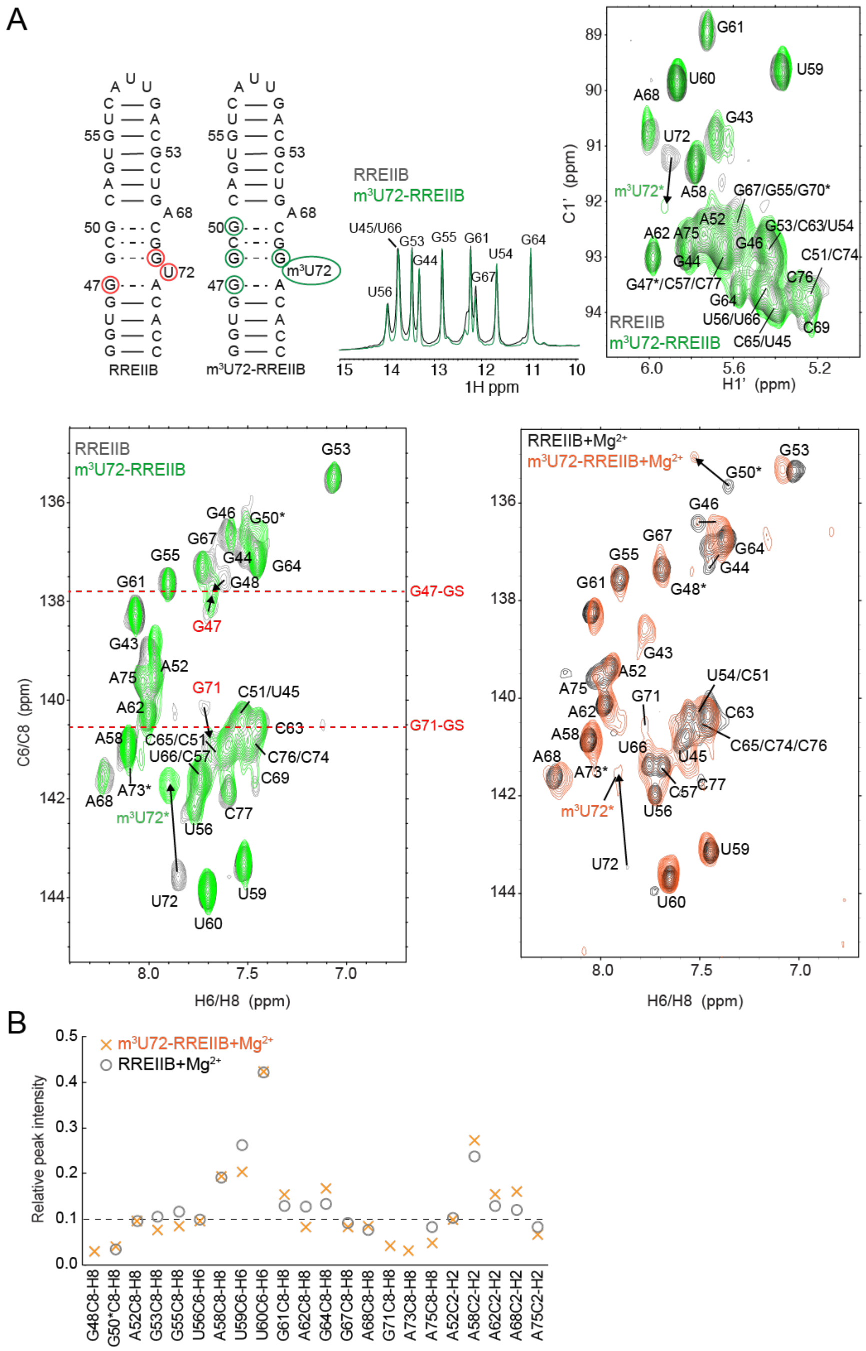


**Figure S7.** NMR spectra and assignments for ES knockout mutant m$^3$U72-RREIIB measured at 25 ºC. (A) The secondary structure of RREIIB and m$^3$U72-RREIIB based on NMR data. Residues showing significant RDs ($p_B$ > 15%, $\Delta\omega$ > 1.5 ppm) or larger differences in chemical shifts between m$^3$U72-RREIIB and RREIIB are highlighted using red circles on RREIIB and green circles on m$^3$U72-RREIIB respectively. Resonances with significant RD (outside from the modified residue) are expected to show larger perturbations toward GS due to suppression of ESs in $m^3$U72-RREIIB. Indeed, G71-C8H8 and G47-C8H8 which have large RD, show large changes (~ 0.3 ppm) toward the GS chemical shift measured by RD in the absence of $Mg^{2+}$ (red dashed lines). U72 experiences perturbations due to the modification. Chemical shift perturbations at G48-C8H8 and G50-C8H8 (+$Mg^{2+}$) which have no RDs could result from slight changes in the GS with introduction of the modification. (B) Normalized resonance intensities measured for RREIIB and $m^3$U72-RREIIB in the presence of 3 mM $Mg^{2+}$. A52-C8H8, A52-C2H2 and U56-C6H6 were used as a reference and normalized to 0.1. G48, G71 and A73, which experience line-broadening due to exchange with ES, show increased intensities in $m^3$U72-RREIIB consistent with suppression of the exchange. Buffer conditions: 15 mM sodium phosphate, 25 mM NaCl, 0.1 mM EDTA, pH 6.4 with or without 3 mM $MgCl_2$.

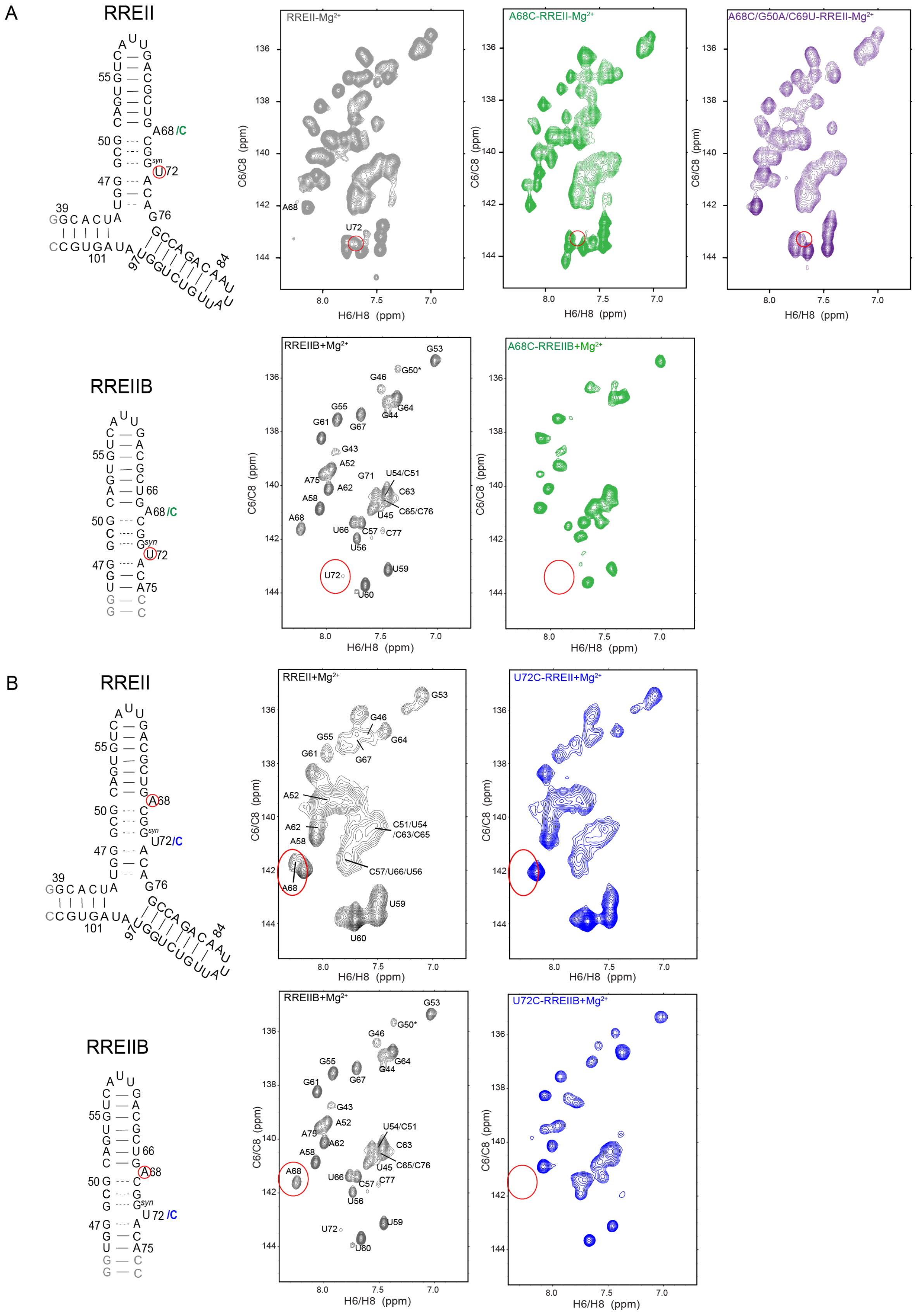
A
RREII
A68/C
U72
RREII-Mg2+
A68C-RREII-Mg2+
A68C/G50A/C69U-RREII-Mg2+
C6/C8 (ppm)
H6/H8 (ppm)
RREIIB
RREIIB+Mg2+
A68C-RREIIB+Mg2+
G53
G50*
G46
G64
G44
G55
G61
G67
G43
A52
G71
U54/C51
C63
A75
A62
A58
C65/C76
U45
A68
U66
C57
C77
U56
U59
U72
U60
B
RREII
A68
U72/C
RREII+Mg2+
U72C-RREII+Mg2+
C51/U54
/C63/C65
C57/U66/U56
RREIIB
U72C-RREIIB+Mg2+

C

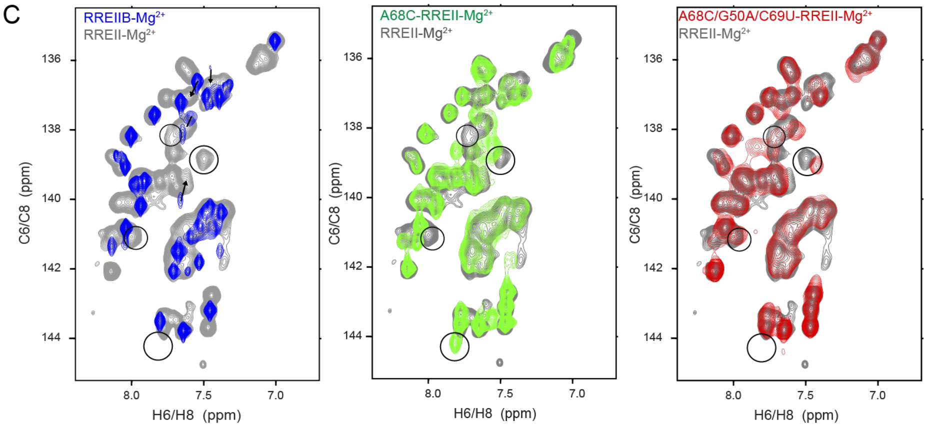


**Figure S8.** NMR spectra of RREII and RREIIB mutants used in fluorescence binding studies measured at 25 ºC. (A, B) Residues that experience broadening due to micro-to-millisecond exchange are highlighted using red circles on the secondary structure. (A) In both ES2-stabalizing A68C-RREII and A68C-RREIIB, U72-C6 was significantly broadened (highlighted in a circle), consistent with a bias toward ES2, and this broadening was partially rescued in the A68C/G50A/C69U-RREII recue mutant. (B) In both ES-stabilizing U72C-RREII and U72C-RREIIB, A68-C8 was significantly broadened (highlighted in a circle) consistent with a bias toward the ES1 and ES2. (C) Spectra showing that the ES-stabilizing mutation A68C induces long-range perturbations (highlighted in circles) at nucleotides remote from the stem IIB. In order to compare potentially dynamic residues, NMR spectra in the absence of $Mg^{2+}$ were compared to avoid further line broadening. Buffer conditions: 15 mM sodium phosphate, 25 mM NaCl, 0.1 mM EDTA, pH 6.4 with or without 3 mM $MgCl_2$.

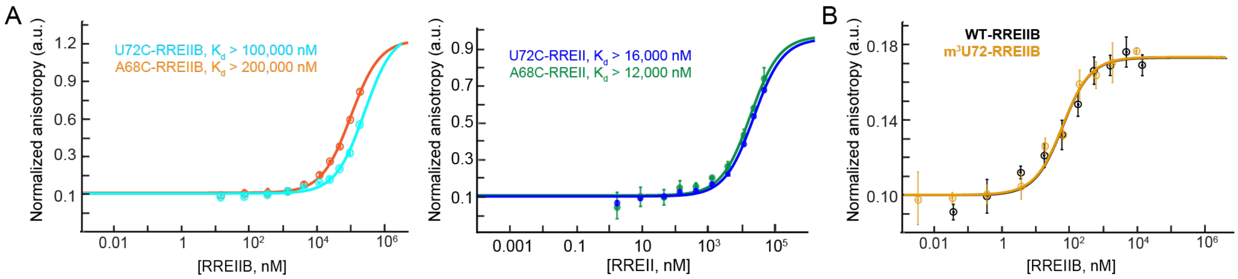


**Figure S9.** Measurement of the binding affinity between Rev-ARM peptide and ES-stabilizing mutants, and ES knockout mutant using fluorescence polarization. (A) Normalized anisotropy values measured for ES stabilizing mutants with higher RRE mutant concentrations using with one-site binding model (see methods). Due to the lower binding affinity, the binding curves did not reach saturation, and further addition of ES-stabilizing mutants resulted in additional low-affinity binding. (B) Normalized anisotropy values measured for RREIIB and $m^3$U72-RREIIB ES knockout mutant fitted with one-site binding model (see methods). The anisotropy value observed in the absence of RNA was normalized to 0.1. Uncertainty reflects the standard deviation from three independent measurements. Buffer conditions: 30 mM HEPES, pH= 7.0, 100 mM KCl, 10mM sodium phosphate, 10 mM ammonium acetate, 10 mM guanidinium chloride, 2 mM $MgCl_2$, 20 mM NaCl, 0.5 mM EDTA, and 0.001% (v/v) Triton-X100. Concentration of Rev-Fl peptide was 10 nM and 1 nM for RREIIB and RREII respectively.

**Table S1**. Exchange parameters obtained from fitting $R_{1\rho}$ data measured on RREIIB.

| *Nuclei* | $\Delta\omega$*_ES1* *(ppm)* | ***pop_ES1*** *(%)* | $k_{ex\_ES1}$ *($s^{-1}$)* | $\Delta\omega$*_ES2* *(ppm)* | ***pop_ES2*** *(%)* | $k_{ex\_ES2}$ *($s^{-1}$)* | $R_1$ ($s^{-1}$) | $R_2$ ($s^{-1}$) | **Red.** $\chi^2$ |
|---|---|---|---|---|---|---|---|---|---|
| **+$Mg^{2+}$** | | | | | | | | | |
| U72-C8 | -2.3± 0.1 | 17.5 ± 2.1 | 6195 ± 423 | -1.2± 0.5 | 3.9 ± 0.1 | 2364 ± 53 | 0.0± 0.6 | 60.4± 3.6 | 0.28 |
| G46-C8 | 1.1± 0.1 | | | 1.1± 0.1 | | | 0.8± 0.3 | 67.6± 1.3 | |
| G67-C8 | | | | 0.9± 0.02 | | | 1.0 ± 0.1 | 65.7± 0.3 | |
| A68-C8 | | | | -2.5± 0.03 | | | 0.8 ± 0.2 | 45.2± 0.4 | |
| **-$Mg^{2+}$** | | | | | | | | | |
| U72-C8 | -2.6± 0.1 | 5.7± 0.4 | 10751± 442 | -0.5± 0.04 | 11.0± 0.4 | 2311± 109 | 0.7± 0.1 | 48.5± 1.1 | 0.45 |
| G71-C8 | -3.8± 0.2 | | | -1.6± 0.05 | | | 0.2± 0.7 | 99.0± 3.7 | |
| G47-C8 | 4.0± 0.2 | | | 1.6± 0.05 | | | 1.0± 0.5 | 51.7± 2.9 | |
| G46-C8 | 1.3± 0.1 | | | 0.7± 0.02 | | | 0.5± 0.1 | 48.8± 1.0 | |
| G67-C8 | | | | 0.4± 0.02 | | | 0.7± 0.2 | 54.6± 0.5 | |
| A68-C8 | | | | -2.4± 0.07 | | | 0.0± 1.2 | 53.7± 2.6 | |

**Table S2:** List of spin-lock powers and offsets used in the $R_{1\rho}$ experiments at 25 ºC. Shown are on-resonance spin-lock power ($\omega 2\pi^{-1}$) or off-resonance spin-lock ($\omega 2\pi^{-1}$) at variable offsets ($\Omega 2\pi^{-1}$) indicated as "$\omega 2\pi^{-1}$ & $\pm$ [$\Omega 2\pi^{-1}$]"

| RREIIB with 3mM $Mg^{2+}$ | On-resonance spinlock power (Hz)/Off-resonance spinlock power (Hz) with [offsets] |
|---|---|
| U72-C6 | 100, 150, 200, 250, 300, 400, 500, 700, 1000, 1200, 1600, 2000, 2400, 2800 |
| | 200 & [-600] ±[50, 100, 150, 200, 250, 300, 400, 500] |
| | 400 & [900, -800] ±[50, 100, 200, 300, 400, 500, 600, 700, 800, 1000, 1100, 1300] |
| | 700 & ±[100, 200, 300, 400, 500, 700, 900, 1100, 1300, 1500, 1700] |
| | 1000 & ±[100, 200, 300, 400, 500, 700, 900, 1100, 1300, 1500, 1700, 1900, 2200] |
| G46-C8 | 100, 150, 200, 250, 300, 400, 500, 700, 900, 1200, 1600, 2000, 2400, 2800 |
| | 150 & ±[50, 100, 150, 200, 250, 300, 350, 400, 450] |
| | 300 & ±[50, 100, 150, 200, 250, 300, 400, 500, 600, 750, 900] |
| | 500 & ±[100, 200, 300, 400, 500, 600, 700, 800, 900, 1100, 1300] |
| | 700 & ±[100, 200, 300, 400, 500, 600, 700, 800, 900, 1100, 1300, 1500, 1700] |
| A68-C8 | 100, 150, 200, 250, 300, 400, 500, 700, 900, 1200, 1600, 2000, 2400, 2800 |
| | 200 & [-600] ±[50, 100, 150, 200, 250, 300, 400, 500] |
| | 400 & [900, -800] ±[50, 100, 200, 300, 400, 500, 600, 700, 800, 1000, 1100, 1300] |
| | 700 & ±[100, 200, 300, 400, 500, 700, 900, 1100, 1300, 1500, 1700] |
| | 1000 & ±[100, 200, 300, 400, 500, 700, 900, 1100, 1300, 1500, 1700, 1900, 2200] |
| G67-C8 | 100, 150, 200, 250, 300, 400, 500, 700, 900, 1200, 1600, 2000, 2400, 2800 |
| | 150 & ±[50, 100, 150, 200, 250, 300, 350, 400, 450] |
| | 300 & ±[50, 100, 150, 200, 250, 300, 400, 500, 600, 750, 900] |
| | 500 & ±[100, 200, 300, 400, 500, 600, 700, 800, 900, 1100, 1300] |
| | 700 & ±[100, 200, 300, 400, 500, 600, 700, 800, 900, 1100, 1300, 1500, 1700] |
| A75-C8 | 250, 400, 500, 700, 900, 1200, 1600, 2000, 2400, 2800 |
| | 400 & ±[100, 200, 300, 400, 500, 600, 700, 900, 1100] |
| | 700 & ±[100, 200, 300, 400, 500, 600, 700, 800, 900, 1100, 1300, 1500, 1700] |
| | 1000 & ±[100, 200, 300, 400, 500, 600, 700, 800, 900, 1100, 1300, 1500, 1700, 2000, 2300, 2600] |
| A52-C8 | 150, 200, 250, 300, 400, 500, 700, 900, 1200, 1600, 2000, 2400, 2800 |
| | 200 & ±[50, 100, 150, 200, 250, 300, 400, 500, 600] |
| | 400 & ±[100, 200, 300, 400, 500, 600, 700, 900, 1100] |
| | 700 & ±[100, 200, 300, 400, 500, 600, 700, 800, 900, 1100, 1300, 1500, 1700] |
| | 1000 & ±[100, 200, 300, 400, 500, 600, 700, 800, 900, 1100, 1300, 1500, 1700, 2000, 2300, 2600] |
| U66-C6 | 150, 200, 250, 300, 400, 500, 700, 1000, 1200, 1600, 2000, 2400, 2800 |
| | 250 & ±[50, 100, 150, 200, 250, 300, 400, 500, 600] |
| | 400 & ±[100, 200, 300, 400, 500, 600, 700, 900, 1100, 1300] |

| | |
|---|---|
| | 700 & ±[100, 200, 300, 400, 500, 700, 900, 1100, 1300, 1500, 1700] |
| | 1000 & ±[100, 200, 300, 400, 500, 700, 900, 1100, 1300, 1500, 1700, 1900, 2200] |
| G53-C8 | 150, 200, 250, 300, 400, 500, 700, 900, 1200, 1600, 2000, 2400, 2800 |
| | 200 & ±[50, 100, 150, 200, 250, 300, 400, 500, 600] |
| | 400 & ±[100, 200, 300, 400, 500, 600, 700, 900, 1100] |
| | 700 & ±[100, 200, 300, 400, 500, 600, 700, 800, 900, 1100, 1300, 1500, 1700] |
| | 1000 & ±[100, 200, 300, 400, 500, 600, 700, 800, 900, 1100, 1300, 1500, 1700, 2000, 2300, 2600] |

| RREIIB without $Mg^{2+}$ | On-resonance spinlock power (Hz)/Off-resonance spinlock power (Hz) with [offsets] |
|---|---|
| U72-C6 | 100, 150, 200, 250, 300, 400, 500, 700, 900, 1200, 1600, 2000, 2400, 2800 |
| | 200 & ±[50, 100, 150, 200, 250, 300, 400, 500, 600] |
| | 500 & ±[100, 200, 300, 400, 500, 600, 700, 900, 1100, 1400] |
| | 800 & ±[100, 200, 300, 400, 500, 600, 700, 900, 1100, 1300, 1600, 1900, 2200] |
| | 1200 & ±[100, 200, 300, 400, 500, 600, 800, 1100, 1400, 1800, 2200, 2700, 3200] |
| | 1600 & ±[100, 200, 300, 500, 700, 900, 1200, 1600, 2000, 2400, 2800, 3200, 3600, 4000] |
| G71-C8 | 150, 200, 250, 300, 400, 500, 700, 900, 1200, 1600, 2000, 2400, 2800 |
| | 200 & ±[50, 100, 150, 200, 250, 300, 400, 500, 600] |
| | 400 & ±[100, 200, 300, 400, 500, 600, 700, 900, 1100] |
| | 700 & ±[100, 200, 300, 400, 500, 600, 700, 800, 900, 1100, 1300, 1500, 1700] |
| | 1000 & ±[100, 200, 300, 400, 500, 600, 700, 800, 900, 1100, 1300, 1500, 1700, 2000, 2300, 2600] |
| G47-C8 | 150, 200, 250, 300, 400, 500, 700, 900, 1200, 1600, 2000, 2400, 2800 |
| | 400 & ±[100, 200, 300, 400, 500, 600, 700, 900, 1100] |
| | 700 & ±[100, 200, 300, 400, 500, 600, 700, 800, 900, 1100, 1300, 1500, 1700] |
| | 1000 & ±[100, 200, 300, 400, 500, 600, 700, 800, 900, 1100, 1300, 1500, 1700, 2000, 2300, 2600] |
| G46-C8 | 100, 150, 200, 250, 300, 400, 500, 700, 900, 1200, 1600, 2000, 2400, 2800 |
| | 150 & ±[50, 100, 150, 200, 250, 300, 350, 400] |
| | 200 & ±[50, 100, 150, 200, 250, 300, 400, 500, 600] |
| | 300 & ±[100, 200, 300, 400, 500, 600, 700, 900] |
| | 400 & ±[100, 200, 300, 400, 500, 600, 700, 900, 1100] |
| | 700 & ±[100, 200, 300, 400, 500, 600, 700, 800, 900, 1100, 1300, 1500, 1700] |
| A68-C8 | 100, 150, 200, 250, 300, 400, 500, 700, 900, 1200, 1600, 2000, 2400, 2800 |
| | 150 & ±[50, 100, 150, 200, 250, 300, 350, 400] |
| | 200 & ±[50, 100, 150, 200, 250, 300, 400, 500, 600] |
| | 300 & ±[100, 200, 300, 400, 500, 600, 700, 800, 900] |
| | 400 & ±[100, 200, 300, 400, 500, 600, 700, 900, 1100] |
| | 700 & ±[100, 200, 300, 400, 500, 600, 700, 800, 900, 1100, 1300, 1500, 1700] |
| G67-C8 | 100, 150, 200, 250, 300, 500, 700, 900, 1200, 1600, 2000, 2400, 2800 |

| | |
|---|---|
| | 200 & ±[50, 100, 150, 200, 250, 300, 400, 500, 600] |
| | 300 & ±[100, 200, 300, 400, 500, 600, 700, 800, 900] |
| | 400 & ±[100, 200, 300, 400, 500, 600, 700, 900, 1100] |
| | 700 & ±[100, 200, 300, 400, 500, 600, 700, 800, 900, 1100, 1300, 1500, 1700] |
| A75-C8 | 150, 200, 250, 300, 400, 500, 700, 900, 1200, 1600, 2000, 2400, 2800 |
| | 200 & ±[50, 100, 150, 200, 250, 300, 400, 500, 600] |
| | 400 & ±[100, 200, 300, 400, 500, 600, 700, 900, 1100] |
| | 700 & ±[100, 200, 300, 400, 500, 600, 700, 800, 900, 1100, 1300, 1500, 1700] |
| | 1000 & ±[100, 200, 300, 400, 500, 600, 700, 800, 900, 1100, 1300, 1500, 1700, 2000, 2300, 2600] |
| A52-C8 | 150, 200, 250, 300, 400, 500, 700, 900, 1200, 1600, 2000, 2400, 2800 |
| | 200 & ±[50, 100, 150, 200, 250, 300, 400, 500, 600] |
| | 400 & ±[100, 200, 300, 400, 500, 600, 700, 900, 1100] |
| | 700 & ±[100, 200, 300, 400, 500, 600, 700, 800, 900, 1100, 1300, 1500, 1700] |
| | 1000 & ±[100, 200, 300, 400, 500, 600, 700, 800, 900, 1100, 1300, 1500, 1700, 2000, 2300, 2600] |

| **RREIIB at pH 8** | **On-resonance spinlock power (Hz)/Off-resonance spinlock power (Hz) with [offsets]** |
|---|---|
| G47-C8 | 100, 150, 200, 250, 300, 400, 500, 700, 900, 1200, 1600, 2000, 2400, 2800 |
| | 200 & ±[50, 100, 150, 200, 250, 300, 400, 500, 600] |
| | 600 & [±100, ±200, ±300, ±400, ±500, ±600, ±700, ±900, ±1100, ±1400, ±1700] |
| | 1000 & [±100, ±200, ±300, ±400, ±500, ±600, ±800, ±1100, ±1400, ±1800, ±2200, ±2700, ±3200] |
| | 1600 & [±100, ±200, ±300, ±500, ±700, ±900, ±1200, ±1600, ±2000, ±2400, ±2800, ±3200, ±3600, ±4000] |
| A68-C8 | 100, 150, 200, 250, 300, 400, 500, 700, 900, 1200, 1600, 2000, 2400, 2800 |
| | 200 & ±[50, 100, 150, 200, 250, 300, 400, 500, 600] |
| | 600 & [±100, ±200, ±300, ±400, ±500, ±600, ±700, ±900, ±1100, ±1400, ±1700] |
| | 1000 & [±100, ±200, ±300, ±400, ±500, ±600, ±800, ±1100, ±1400, ±1800, ±2200, ±2700, ±3200] |
| | 1600 & [±100, ±200, ±300, ±500, ±700, ±900, ±1200, ±1600, ±2000, ±2400, ±2800, ±3200, ±3600, ±4000] |
